# MHConstructor: A high-throughput, haplotype-informed solution to the MHC assembly challenge

**DOI:** 10.1101/2024.05.20.595060

**Authors:** Kristen J. Wade, Rayo Suseno, Kerry Kizer, Jacqueline Williams, Juliano Boquett, Stacy Caillier, Nicholas R. Pollock, Adam Renschen, Adam Santaniello, Jorge R. Oksenberg, Paul J. Norman, Danillo G. Augusto, Jill A. Hollenbach

## Abstract

The extremely high levels of genetic polymorphism within the human major histocompatibility complex (MHC) limit the usefulness of reference-based alignment methods for sequence assembly. We incorporate a short read *de novo* assembly algorithm into a workflow for novel application to the MHC. MHConstructor is a containerized pipeline designed for high-throughput, haplotype-informed, reproducible assembly of both whole genome sequencing and target-capture short read data in large, population cohorts. To-date, no other self-contained tool exists for the generation of *de novo* MHC assemblies from short read data. MHConstructor facilitates wide-spread access to high quality, alignment-free MHC sequence analysis.

## Background

The vast majority of human diseases have a complex, polygenic component, with risk loci often spread throughout the genome (Boyle *et al*. 2017). Notably, the single region of the human genome with the greatest number of trait and disease association signals is the major histocompatibility complex (MHC) (Lenz et al., 2016). The MHC is located on the short arm of chromosome 6, at 6p21.31. MHC genomic variation is strongly associated with its role in the adaptive immune system primarily due to the presence of the human leukocyte antigen (HLA) genes. HLA class I and class II genes encode the antigen-presenting, surface-marker proteins responsible for driving the adaptive immune responses. Over evolutionary time, this critical immune function has been subjected to a diversity of selective scenarios, ultimately producing the single most polymorphic region of the human genome (Doherty and Zinkernagel 1975; Hudson and Kaplan 1988; Wakeland *et al*. 1990; Takahata and Satta 1998; Spurgin and Richardson 2010; Robinson *et al*. 2017; Kaufman 2018; Radwan *et al*. 2020; Talarico *et al*. 2021). While most disease association studies have focused on variation in HLA, the extended MHC region contains over 165 protein-coding gene, many involved in the immune response (Trowsdale and Knight 2013).

The immense genetic diversity critical for immune responses has made it difficult to thoroughly elucidate the molecular mechanisms by which MHC variation contributes to numerous diseases. Advances toward this end have been primarily driven by targeted analysis of protein-coding genes or genome-wide association studies (GWAS) conducted with single nucleotide polymorphism (SNP) arrays (Beecham *et al*. 2013, 2022; Hollenbach and Oksenberg 2015; Lenz *et al*. 2016; Matzaraki *et al*. 2017). While many disease associated SNPs map to the classical HLA genes, many HLA-independent associations throughout the MHC have also been identified. However, our current understanding of the role of MHC variation in disease remains limited since this region is often excluded from whole genome analysis, which provides greater resolution variant maps than SNP arrays. This is not for lack of interest in examining high-resolution nucleotide variation, but due to the significant challenges inherent to a robust assembly of short read data across the 5Mbp MHC region. In particular, novel heterozygosity, copy number variation, and large structural rearrangements are challenging to identify using methods that merely align sequencing reads to a reference sequence (Dilthey 2021).

*De novo* assembly offers an appealing solution to capture variation in a reference-free manner.(Khan *et al*. 2018). The MHC sequences associated with the 1000GenomesProject phase 3 data are composed of high-quality, 30x coverage, short-read NGS data that has been *de novo* assembled (Auton *et al*. 2015). A consortia in Denmark released 100 *de novo* assembled MHC sequences from family trio haplotypes to serve as population references (Maretty *et al*. 2017). That same year, 95 novel extended MHC sequences derived from human cell lines were assembled *de novo* using NGS short-read sequencing (Norman *et al*. 2017). These assemblies successfully recreated known structural variation and identified extensive novel variation. Long-read sequencing is another promising solution to the challenges of highly heterozygous diploid assembly. Recently, Houwaart et al. performed a follow-up study in seven, consanguineous, HLA-homozygous cell lines from Norman et al., 2017, representing the major HLA class II and C4 haplotype categories observed so far. Through a combination of Illumina short read and HiFi long read sequencing, they were able to fully characterize these seven MHC sequences at unprecedented, highly accurate single nucleotide resolution (Houwaart et al., 2023). This work contributed greatly needed high-quality reference haplotype sequences. Although long-read technology is becoming the industry standard for high-quality reference genomes (Rhie *et al*. 2021; Houwaart *et al*. 2023), its applicability to population-level research, such as large (n>=500) disease association and demographic studies, remains resource-prohibitive. Similarly, trio-phasing using haplotype-resolved assemblies has proven an effective, but expensive, strategy for handling the complexities of diploid MHC assembly (Jensen *et al*. 2017; Koren *et al*. 2018). Trio binning of short-read data can be used to separate haplotype-unique long-read content from each parent (Koren *et al*. 2018). Binned sequencing data is then assembled separately, *de novo*, generating two highly accurate, haplotype-resolved haploid consensus sequences per individual, one for each parental chromosome. While trio cohorts are ideal, the tripled sequencing cost and assembly time is rarely feasible for large disease cohorts. Further, the collection of family data in disease contexts can be very difficult, especially in non-pediatric diseases.

Here, we implement an MHC reference haplotype-guided, *de novo* assembly approach, based on an assembly process originally designed to construct novel genomes when no species reference genome is available (Lischer and Shimizu 2017). To do this, we establish the class II HLA –DR haplotype and C4 status for each individual. Since these regions represent the major classes of known, large, structural variation observed across the extended MHC (Houwaart *et al.,* 2023), we use these to define ‘MHC haplotypes’. In this way, we simulate trio binning to handle heterozygosity in the large structural variation associated with the HLA class II and C4 regions, while still capturing the remaining MHC sequence variation in a reference-free manner. Further, while many advances have been made in handling the problem of MHC sequence assembly, to-date, none have been developed into a self-contained tool that can be used reproducibly or that can scale to large, population level cohorts. The large number of software and associated dependencies required to perform *de novo* assembly alone, often creates a barrier to accessibility and reproducibility, particularly as version control is widely inconsistent across bioinformatic tools (Baker 2016; Cohen-Boulakia *et al*. 2017; Cokelaer *et al*. 2023). These challenges have resulted in limited capacity to reliably generate MHC sequences and thoroughly query the functional role of MHC variation in disease association contexts. We here present MHConstructor, a Singularity container (Kurtzer *et al*. 2017) which houses an integrated suite of tools aimed at producing high-quality MHC sequence at high-throughput: A *de novo* assembly pipeline (Lischer and Shimizu 2017) custom adapted for application to the extended 5Mb MHC region, a short read quality control filter (Lischer and Shimizu 2017), *C4Investigator* for complement component 4 (C4) genotyping and copy number variation calling (Marin *et al*. 2024), *T1K* for first field DRB1 genotyping (Song *et al*. 2023), a batch short read data extraction that interfaces with the 1000GenomesProject database (Auton *et al*. 2015), and all necessary supplementary software dependencies, in a user-friendly, containerized interface.

Recent work has demonstrated that even with all that is currently known of MHC diversity, it remains an underestimation of the true extent of polymorphism within this complex genome region. By developing and implementing the high-throughput, haplotype-informed *de novo* assembly method MHConstructor, we provide the means to apply a validated pipeline to the hundreds of existing disease cohorts analyzed with short-read NGS and those newly created. We also generate an additional 536 novel MHC haplotype assemblies from 369 individuals. In doing so, we increase access to a fuller spectrum of MHC diversity, which will improve our understanding of how variation in this critical region contributes to human complex genetic disease.

## Results

### 1. Haplotype-guided, *de novo* short read MHC assembly workflow

We build from current short-read *de novo* assembly progress and develop a tool designed specifically for *MHC de novo* assembly. We simulate trio-binning by substituting parental sequence data with inferred, proxy parental MHC sequences, from among the eight major MHC reference sequences (Houwaart *et al*. 2023). We describe these as “MHC haplotypes” (Supplemental Documents 2 and 3). These proxy-parental MHC haplotypes are then used as the guide sequences to facilitate haplotype-guided, *de novo* assembly. This strategy provides a convenient framework for handling heterozygous, diploid sequencing data in a haplotype-informed manner. We implement and adapt a previously established pipeline, which was originally designed in a comparative species context, to capture novel, species-specific sequence when a reference genome was not available for alignment (Lischer and Shimizu 2017). We then carry out the *de novo* assembly process in a haplotype-informed manner, generating pseudo-haploid assemblies, rather than a combined diploid assembly, for each individual (Figure 1).

**Figure 1.**
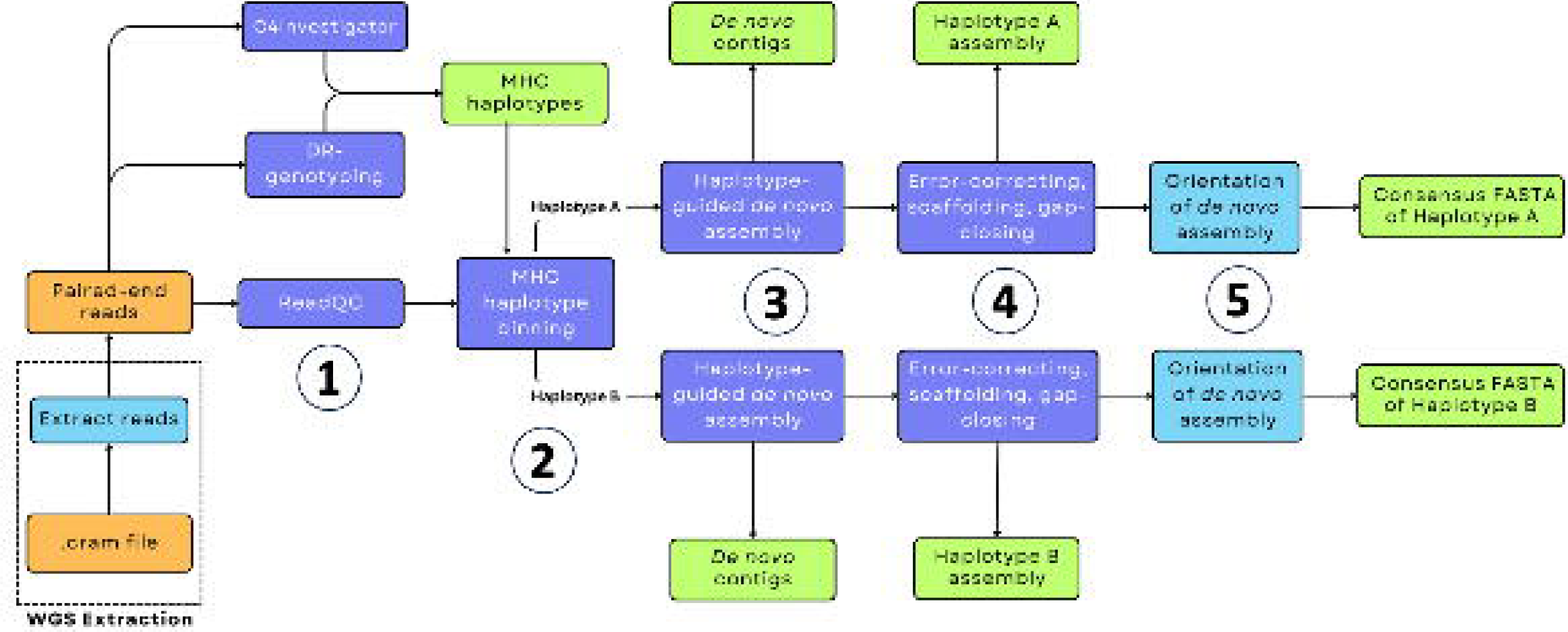
MHConstructor workflow for haplotype-informed, *de novo* MHC assembly. Orange indicates source data. Purple indicates previously published software. Teal indicates novel software. Green indicates output data.

### 2. Evaluation of assembly performance metrics

#### 2.1 Assembly speed and quality are impacted by assembly kmer size and sequencing depth

*De novo* assembly algorithms are inherently computationally intensive, which can introduce implementation challenges in a high throughput context. To characterize how our algorithm responds to various parametric spaces, we have tested assembly parameters on a high-performance computing cluster and describe the relationship of key assembly parameters, assembly kmer size and sequencing depth to overall performance. Small kmer sizes are considered more successful at high accuracy mapping, whereas longer kmer sizes can improve repeat content assembly. To generate an initial range of starting kmer sizes to evaluate, kmerGenie was applied to a set of n=20 samples, assessing the mapped reads vs unmapped reads separately. Higher kmer values were found to be more relevant for reads which were able to be mapped to the haplotype guide sequence (k=41-61) (Supplemental Table S1), while lower sizes of k were predicted for the unmapped reads (k=21-26) (Supplemental Table S2). Kmer size predictions for unmappable reads showed less variation between individuals than the mappable reads. For reads which mapped to the guide haplotype, best kmer size varied significantly across samples (stdev 4.15-21.19). Additionally, the number of reads evaluated impacts the best k-mer size identified, with greater sequencing depth allowing for larger kmer sizes. However, variability in predicted best kmer size between samples also increases with greater sequencing depth. Overall, there is less variability in kmer prediction for reads which do not initially map to guide haplotype, and these values are generally lower than those predicted for the well mapped reads, consistently falling between k=21-26 for all sequencing depths considered (Supplemental Table S2).

We also empirically tested *de novo* assembly kmer size by evaluating assembly quality metrics for varying kmer sizes and find that using larger kmer sizes for mapped reads produce stronger assembly metrics (Supplemental Figure S1). Additionally, we measured how assembly speed is impacted by choice of kmer size. When the kmer size used to assemble unmapped reads is small (<30), user time for the entire analysis increases in an approximately exponential scale with respect to starting read count (Supplemental Table S3). However, at k=51, for both unmapped reads and mapped reads the relationship between starting read count and analysis time becomes linear (Supplemental Table S4). For this reason, though Kmergenie predicted a smaller kmer size to be most accurate for the unmappable reads, we chose to use larger values in the interest of speed. Using the above metrics, we chose k=51 for *de novo* assembly of both haplotype-mapping and non-mapping reads. Unsurprisingly, we find that better assembly metrics are correlated with higher sequencing depth (Supplemental Figure S2). However, we do find that total length of *de novo* assembled contig sequence generated does reach an upper limit at around ∼4.6Mb. This may be indicative of the upper limit of sequence that can be *de novo* assembled with short read data. We find that for target-capture samples with an average sequencing depth of 60X, the full analysis takes approximately one and a half hours to complete per MHC haplotype, with multithreading capacity (Table 1). The WGS 30X 1KGP samples, however, were much quicker to assemble, with an average time of 26 minutes per MHC haplotype (Table 1).

**Table 1.** Average runtime (in user minutes) for MHConstructor assembly, at eight threads on high performance computing cluster.

| Sequencing depth | *de novo* assembly<br>(min) | Error correction+scaffolding<br>(min) | Total assembly<br>run (min) |
| --- | --- | --- | --- |
| 65X target capture | 53 | 51 | 104 |
| 30X WGS | 13 | 13 | 26 |

### 3. Quantification of MHConstructor error rates using high-quality MHC reference sequences

#### 3.1 Haplotype-informed assembly validation via re-creation of known, fully characterized MHC sequences

In addition to quantifying overall assembly quality metrics, we also described the error rate inherent to the short read assembly process. We first evaluated this using the set of high quality references representing haploid MHC sequences which have also been incorporated into MHConstructor as guide sequences (Houwaart *et al*. 2023). Using the Illumina short read dataset associated with each of the haploid MHC reference sequences (*SRA:* SRP348947, *BioProject:* <u>PRJNA764575</u>), we re-assembled each sequence *de novo* using MHConstructor. Summary statistics for these assemblies can be found in Supplemental Table S5. Reference MHC coverage was incomplete (Supplemental Tables S6,S7). However, based on our previous findings, we inferred that this was due to the low starting read count (Supplementary Table S5), corresponding to a <30X average coverage across the MHC, limiting our ability to recover the full region. Therefore, we calculated the error statistics as a percent of the total assembled sequence, rather than as a percentage of the entire length of the MHC. We found that the average percent error for basecalling (base call error %) was 0.031%, the average error attributable to non-repeat associated false SVs (>1bp) and the average error attributable to repeat-derived false SVs was 0.05% of the total, assembled sequence for each haplotype (Table 2). We also evaluated whether these assemblies were able to construct the correct *HLA* alleles for these haplotypes and found that BLAST alignments revealed that the correct, known alleles corresponded to the strongest percent identity allele match (Supplementary Tables S8 and S9).

**Table 2.** Target-capture MHC assemblies exhibit 0.119% average total error in.

| MHC haplotype | Total assembly length (bp) | Base call error (%) | Non-repeat associated SV error (%) | Repeat-associated SV error (%) | Total error (%) |
| --- | --- | --- | --- | --- | --- |
| OK649231 | 4127329 | 0.0267 | 0.0350 | 0.0899 | 0.1516 |
| OK649232 | 3908195 | 0.0311 | 0.0515 | 0.1144 | 0.1970 |
| OK649234 | 4539318 | 0.0218 | 0.0441 | 0.0068 | 0.0727 |
| OK649235 | 3710459 | 0.0554 | 0.0215 | 0.0124 | 0.0893 |
| OK649236 | 4196340 | 0.0211 | 0.0528 | 0.0080 | 0.0819 |

SV: structural variant ≥ 1bp in length

#### 3.2. Use of phased, long-read MHC sequences to determine diploid *de novo* assembly error rate

To further validate this method and ensure that it can successfully handle the heterozygous, diploid sequence data found in human populations, we used MHConstructor to re-assemble phased, diploid MHC reference sequences (Huijse *et al*. 2023) from corresponding 1000GenomesProject 30X WGS reads. We generated the assemblies using the long-read sequenced, phased haplotype sequence as the guide sequence. We find that no large structural variants (>1kb) are erroneously introduced (Supplemental Figure S3) when using a threshold on the Ragtag scaffold orientation, which filters out any scaffolds that have a location score <0.1, or location score <0.3 and an orientation score <0.75. In order to evaluate nucleotide composition error, we aligned MHConstructor consensus haplotypes to their long-read, phased haplotypes using *minimap2* to describe base call errors (Li 2018) and *Assemblytics* to identify structural errors (>=1bp INDELs) (Nattestad and Schatz 2016). The amount of mismatched sequence between the two was determined by counting the number of variants identified and dividing by the total length of the MHConstructor consensus haplotype. We found that the average percent error for base calling (base call error %) was 0.17%, the average error attributable to non-repeat associated false SVs (>1bp) was 0.18% and the average error attributable to repeat-derived false SVs was 0.1% of the total, assembled sequence for each haplotype (Table 3). Assessment of the categories and sizes of false structural variants (>1bp) revealed low frequency of repeat expansions and repeat contractions. Across the six MHC haplotypes evaluated, we only found three instances of false structural variants greater then or equal to 500bp in length. Two instances were caused by a 630bp and 646bp repeat expansions, and one was caused by a 2,842 bp repeat contraction (Supplemental Figure S4-S6). Otherwise, all false SVs identified were under 500bp in length (Supplemental Figure S4-S6). We found that of the sites in the MHConstructor assembled haplotypes that did not match the phased, long-read haplotype guide sequence, between 34-54% were a result of true assembly error, with the remaining sites caused by improper inclusion of a heterozygous allele attributed to the other phased haplotype (Supplemental Table S8).

**Table 3.** MHC assemblies from WGS data exhibit an average of 0.44% total error in 1000GenomesProject individuals, SV>=1bp.

| MHC haplotype | Total assembly length (bp) | Base call error (%) | Non-repeat associated SV error (%) | Repeat-associated SV error (%) | Total error (%) |
| --- | --- | --- | --- | --- | --- |
| HG00621.1 | 4684211 | 0.136 | 0.188 | 0.089 | 0.414 |
| HG00621.2 | 4654849 | 0.162 | 0.162 | 0.107 | 0.430 |
| NA19240.1 | 4739094 | 0.179 | 0.162 | 0.065 | 0.407 |
| NA19240.2 | 4728464 | 0.211 | 0.160 | 0.072 | 0.442 |
| NA20129.1 | 4713830 | 0.145 | 0.186 | 0.177 | 0.509 |
| NA20129.2 | 4538560 | 0.162 | 0.195 | 0.058 | 0.415 |

SV: structural variant >= 1bp in length

#### 3.3 HLA class II haplotype-defining structural variation is correctly assembled

The eight primary MHC haplotypes used here as guide sequences are distinguished primarily by structural variation in the HLA class II region. This variation is primarily driven by *HLA-DRB1*, its three duplication paralogs, *HLA-DRB3*, *HLA-DRB4* and *HLA-DRB5*, as well as several pseudogenes. Most individuals have only partial representation of these. The generally observed combinations are *HLA-DRB1* with *HLA-DRB5* (DR2), *HLA-DRB1* with *HLA-DRB3* (DR3) and *HLA-DRB1* with *HLA-DRB4* (DR4) (Rollini *et al*. 1985; Maiers *et al*. 2007). Of these three, the DR4 haplotype contains extra sequence content between *HLA-DRB4* and *HLA-DRB1*, thus extending the length of the MHC and providing a challenge to correct sequence assembly. To ensure that our *de novo* assembly method is capable of properly capturing this structural variation, we assessed the reconstruction of structural variation associated with the HLA-DRB1,-DRB3,-DRB4,-DRB5 genes via dotplot visualization of oriented *de novo* assemblies aligned to the MHC phased reference sequences (Figure 2), using the R package *paftools*. When assemblies are aligned to the haplotype reference with their correctly matched DR-status, there is continuous alignment over the class II region (Figure 2A). Conversely, when aligned to a haplotype reference with a mismatched DR-status (Figure 2B), we see a large break in the alignment, corresponding to the length of the expected structural variation. We find that our approach is capable of correctly assembling this structurally-varied region. This was additionally confirmed through visual inspection in Mauve (Supplemental Figure S7), which we demonstrate with an example from an individual with European ancestry (individual Z) who is heterozygous for DR3 and DR4. Two Individual Z consensus sequences were generated by aligning the *de novo* assembly to the APD (DR3) and DBB (DR4) reference sequences separately. Next, these alignments were visualized in Mauve (Darling *et al*. 2004). The absence of sequence gaps in the alignment shows that the structure was correctly assembled in both cases. Promisingly, we find that even in the event of choosing a mismatched guide MHC for the assembly process (ie; DR4 instead of DR2), the correct class II structure will still be constructed, as long as assembly is scaffolded with respect to the correct, DR-matched guide, though with lower resolution with respect to identification of novel variation (Supplementary Figure S8).

**Figure 2.**
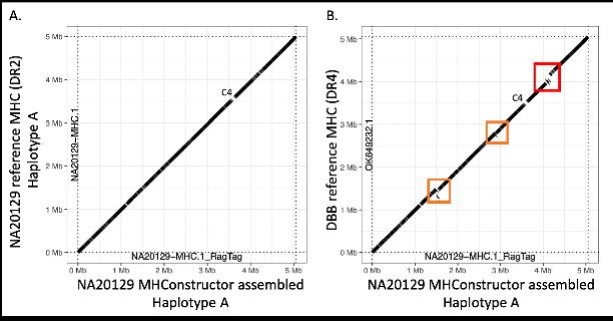
HLA Class II structural variation is correctly assembled. Dotplots depict the MHConstructor, haplotype consensus sequence for 1000GenomesPorject ASW individual NA20129, aligned to A) the correct, phased reference haplotype, which has the short, DR3 HLA class II status and B) to reference haplotype DBB, which has the long, DR4 HLA class II status. Solid diagonal line indicates continuous alignment. Breaks in the line indicate assembly gaps. The red box indicates structural variation resulting from mis-matched HLA class II status. Orange boxes indicate additional regions exhibiting structural variation. The position corresponding to C4 is designated.

#### 3.4 Assembly of short read data reliably describes repetitive content

Since repetitive elements are known to comprise 50-52% of the MHC (Houwaart *et al*., 2023) and roughly 60% of the genome as a whole (deKoning *et al.,* 2011), we wanted to ensure that this method was reliably assembling genomic sequence composed of repetitive content. All final assembled sequences were annotated with RepeatMasker (RM) software, using default settings (www.repeatmasker.org). We then compared our novel assembly RM predictions to RM predictions for the original, phased, MHC reference sequences. We find that repeat element annotations are highly syntenic and differ only by ∼3% of total annotated repeat content (Supplemental Table S9). Further, we verify that our assembly method can properly handle short read data and sensitively assemble the sequences of repetitive elements in the MHC.

### 4. MHConstructor assembly of target-capture 60x and 1000genomes 30x WGS short read MHC data

#### 4.1 Haplotype-guided, *de novo* assembly generates high-quality extended MHC assemblies

MHConstructor was then used to generate 536 novel MHC haplotype assemblies from MHC target capture short read sequencing data. Assembly quality is generally evaluated according to several primary metrics. N50 is a traditional indicator of assembly quality, as it provides a description of contig size distribution, with a larger N50 indicating more contigs of greater length. However, this metric is now recognized by the bioinformatics community to not be wholly representative of assembly quality, especially in genomic regions with complex sequence content. Therefore, we also describe the number of contigs per assembly, as well assembly length (Supplemental Documents 4 and 5). In order to use these scores to meaningfully evaluate the performance of the reference-guided, *de novo* assembly method, we compared them to a previously published dataset of short read, *de novo* assembled MHC sequences, n=95 haplotypes (Norman et al., 2017), based on assembled contigs, prior to scaffolding. For the target-capture assemblies (n= 536 haplotypes), we find that while the distributions show much overlap, t-testing revealed that the target-capture contig number distribution was significantly lower than the distribution from Norman et al. (mean=1774.924 vs. mean=2850.116, p< 2.2e-16), indicative of more complete assemblies. Correspondingly, the target-capture N50 values were significantly larger (mean= 4359.881vs. mean=3894.989, p=9.584e-10), indicating larger, median contig length. There was a slight, but significant, decrease in total assembly length in the target capture compared to the Norman et al. sequences (mean=4185033 vs mean=4567339, p=0.0008554). The same trends were observed in the WGS data, though contig number and N50 value had a completely separate distribution from both the target-capture and the Norman et al. assemblies (Supplemental Figure S9, Supplemental Documents 4 and 5).

#### 4.2. MHConstructor assemblies of target-capture, ASW-like sequences exhibit different assembly metrics than CEU-like sequences

Interestingly, there is a bimodal distribution within the target capture assembly metric distributions (Figure 3). The ASW-like assemblies have a higher contig number (p-value < 2.2e-16), lower N50 (p-value < 2.2e-16) and lower total assembly length (p-value = 8.673e-16) than the CEU-like assemblies. Traditionally, these values are associated with lower assembly quality. However, when comparing assembly metric distributions between the CEU and ASW assemblies generated from 30X WGS data, no differences were observed in contig number (p-value = 0.3184) or N50 length (p-value= 0.2629), though total assembly length was slightly higher in the CEU population (p-value= 0.004869). This difference appears to also be reflected in the pre-assembly short read data, as compared between WGS and target-capture reads. The ratio of unique 150mers in the ASW group relative to the CEU groups, differed between sequencing methods. In the target-capture set, the mean unique 150mers per ASW-like reads was significantly higher than the mean of the CEU-like reads (p-value < 2.2e-16). However, this trend did not occur in the WGS data set (p=0.5212) (Supplemental Figure S10). This trend remained, even following random subsampling of the target-capture individuals to achieve equivalent sample sizes (p-value < 2.2e-16).

**Figure 3.**
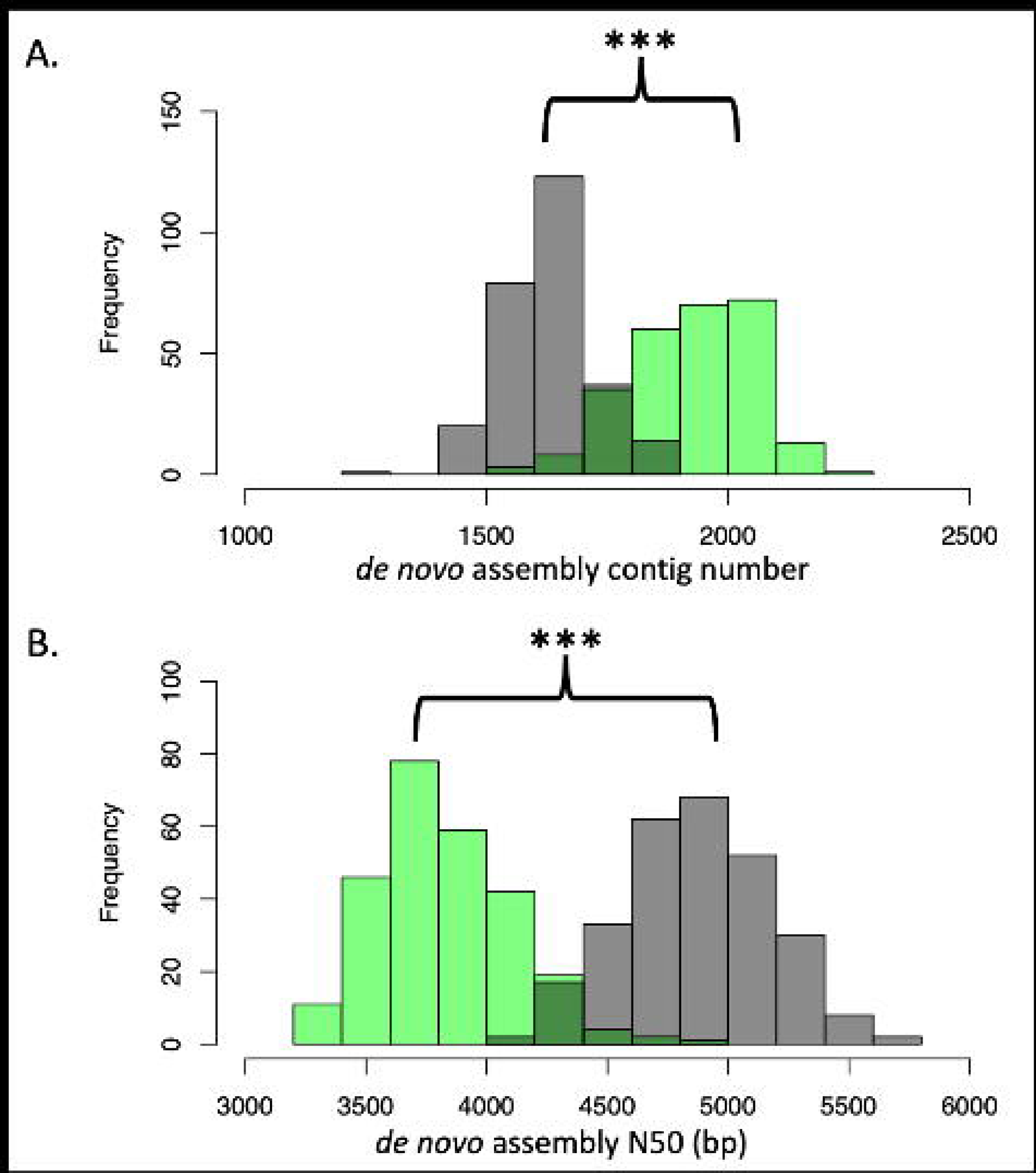
Target-capture assemblies from ASW-like individuals demonstrate different assembly metrics than those from CEU-like individuals. Green distributions represent ASW-like assemblies generated from target-capture, 60X short read data, n=262 MHC haplotypes. Grey distributions represent CEU-like assemblies generated from target-capture, 60X short read data, n=274 MHC haplotypes. A) Histogram of the number of contigs per assembly. B) Histogram of N50 (bp), per assembly.

## Discussion

Though it is known that human MHC variation is implicated in numerous disease phenotypes, aside from the classical *HLA* loci, characterization of variation in this region has been limited to SNP genotyping arrays, reference-based alignment methods, and only more recently, long-read sequencing. Because of this, MHC sequences from hundreds of thousands of disease cohorts with short-read data have remained untapped, and knowledge of the true extent of MHC variation has been obscured. Here, we developed and demonstrated a tool, MHConstructor, that produces high-resolution, *de novo* short read assembly of the extended MHC, for both target-capture and WGS short read sequencing methods.

This tool facilitates enhanced MHC research in three key areas that, to our knowledge, have not yet been attained for high throughput analysis of large, short-read MHC datasets: 1) accuracy, 2) speed, 3) reproducibility and optimization.

1) Accurate sequence assembly. By comparing MHConstructor performance for both target capture and WGS methods, against high quality, reference MHC haplotypes from long-read sequencing data, we have established an error rate between 0.12% and 0.44% overall and minimal error in repetitive regions. We deemed this an acceptable error rate, as the primary use of these sequences will be for the purpose of population-scale, disease association studies. Any spurious variants that occur under this frequency threshold will not be expected to demonstrate high enough frequency at the population level to result in false associations, as most statistical models do not have the sensitivity to detect associations with variants with a population frequency <5%. We also demonstrate that the use of a haplotype-guided *de novo* assembly ensures accurate reconstruction of the known *HLA* Class II structural variation.
2) Rapid assembly speeds: We find that, while speed is impacted to some extent by coverage depth and choice of starting parameters, average runtimes per haplotype range between one hour to one and a half hours, using eight parallel computing threads at computationally intensive stages. Assembly speed can also be impacted by assembly parameters.
3) Containerized and reproducible analysis workflow: Due to the large number of assembly software that these processes rely on, the containerized workflow is essential for establishing this method as a usable tool for a wide variety of audiences. By developing this pipeline structure, executed from within a Singularity container, we have made *de novo* assembly significantly more accessible to a broader range of researchers. Further, this structure ensures that results can be replicated, and facilitates additional optimization and improvement due to its modularity, which allows the user to optimize the analysis parameters for each unique dataset, if desired.

Though MHConstructor generates first field genotyping of *HLA-DRB1* using *T1K* (Song *et al*. 2023), we do not recommend using this data, or the MHConstructor pipeline, for the purposes of clinical *HLA* genotyping. The primary goal of this analysis is to expand access to large population-level cohorts and foster the discovery of novel variation and factors impacting MHC biology across the extended 5 Mbp region. It is not designed to meet the level of stringency required for assigning *HLA* genotypes in a clinical context as there are many robust and user-friendly tools and protocols which are designed for that level of required accuracy, and we encourage users to pursue those, should their interest be in clinical HLA typing.

Throughout the process of our subsampling and optimization analyses, we find that sequencing depths below 25x on average across the MHC are not sufficient to generate full assemblies. We find that assemblies at these lower coverage ranges are more fragmented and do not produce full-length consensus sequences. Therefore, we recommend a sequencing depth of 30x or higher to achieve the most robust assemblies. Higher starting sequencing coverage (45x and 60x) will generally have higher accuracy but will take longer to complete the assembly. There are many additional parameters involved in this analysis which may require fine-tuning and we encourage users to consider these with respect to their own project and data. Since every NGS dataset will have unique quality dynamics and experimental questions, we recommend that users perform a similar evaluation when choosing the most relevant parameters for their dataset.

Though overall assembly statistics appear canonically better in the 30X WGS assemblies, the higher error rate as compared to target-capture sequencing implies that these statistics may be falsely inflated. It is possible that the 30X WGS method is not sensitive enough to fully capture the extent of nucleotide diversity across all individuals. This is additionally supported by the comparison of assembly metrics between the two distinct populations considered. In the target-capture assemblies, we find “lower” quality scores associated with the assemblies of ASW-like individuals, compared to the CEU-like individuals, suggestive of increased nucleotide diversity in the ASW-like population that may be further challenging the *de novo* assembly process. This difference is not observed between the 1KGP 30X WGS ASW and CEU populations, suggesting that the target capture approach may be picking up more of the nucleotide diversity known to be associated with populations of African American ancestry (Campbell and Tishkoff 2008; Tucci and Akey 2019). We find additional support for this interpretation, given that reads from the target-capture sequencing appear capable of identifying a higher amount of likely novel MHC variation in populations with African ancestry than WGS sequencing. It is also likely that the lower, 30X coverage typical of most WGS is performed at may also be contributing to reduced sensitivity. Thus, some novel variation may not even be identifiable with a 30X WGS approach. Though it is unclear if increasing the WGS coverage depth would account for this. Our results suggest this may have an even greater impact in populations with African ancestry, as opposed to European ancestry. It is likely that we may still be underestimating the true extent of MHC variation.

Due to the nature of short-read data, there are certain inherent technical limitations usually described in short-read assemblies. This is primarily evident in regions containing highly repetitive elements. As evidenced by the dotplot visualization showing assembly alignment gaps corresponding to the position of *C4A* and *C4B*, *de novo* short read assembly is not capable of discerning between *C4A* and *C4B* and their duplicates. To avoid this problem, we have implemented *C4Investigator* (Marin *et al*. 2024) annotations, which designate the presence of *C4A* and *C4B*, long and short forms, and copy numbers. Another limitation associated with short read assembly of repeat content is the proper boundary designation and copy number identification of interspersed repeat elements. However, by comparison to long-read assembled, fully phased MHC sequences, we find that our method only introduces an additional 0.086-0.125% error attributable to misassembling interspersed repeats. The other major source of repeat-associated error that short assemblies experience can be caused by incorrect prediction of structural variant breakpoints that may occur within a repetitive sequence. This is an ongoing challenge in the field, and there are many specialized software packages designed to address this single issue (Kosugi *et al*. 2023). Building on our preliminary SV predictions provided using output from MHConstructor, it is likely that future work may further refine these predictions through additional analysis using such algorithms. This will be of importance for disease association studies, as the role of non-coding and repetitive variation in contributing to disease pathogenesis is increasingly becoming recognized (Stankiewicz and Lupski 2010; Madireddy and Gerhardt 2017; Zhou and Zhao 2018; Frydas *et al*. 2022; Liao *et al*. 2023).

### Conclusions

From these haplotype-specific assemblies, variant information can be generated in a variety of ways, including: 1) using an individual’s haplotype-informed consensus sequence(s) to generate variant calls against the standard human reference MHC, hg38. This allows for the identification of haplotype-informed, novel variation, with respect to a single, consistent coordinate system, which is essential for current GWAS models, which rely on input to be consistent across the population. In anticipation of this being directly usable, we have included a method of merging variant calls between two, haplotype-informed consensus sequences into a single variant call file per individual. This can be directly incorporated as input for downstream GWAS tools, such as *plink* (Purcell *et al*. 2007). Alternatively, this data allows for option 2): incorporation of both haplotype-informed consensus sequences into a disease cohort-wide, pan genome graph model, which more robustly captures population-wide variation (Dilthey *et al*. 2015; Ebler *et al*. 2022). These models are still being functionally integrated into GWAS methods, but when efforts reach a widely available format, the sequences output from this analysis can readily be incorporated.

The modular nature of the pipeline workflow lends itself to adaptions and future modifications. Though it has not been tested within the scope of this manuscript, we anticipate that it should be possible to integrate long read sequencing data with this pipeline as well. This could be introduced at Step 3, replacing *de novo* assembled, short read contigs. The use of hybrid long and short-read sequencing is well known to be the most effective method of high-accuracy sequence assembly, and including such data here would likely increase assembly accuracy even further. Future work can be done to fine-tune and validate this alternative application. Additional future efforts will likely involve the application of additional methods to enhance the annotation of SVs in the MHC. It is not a trivial task, and algorithms for the characterization of complicated SVs are continuously being improved upon (Kosugi *et al*. 2023). Future work to formally phase the output of MHConstructor will expand its usability even further (Kajitani *et al*. 2019). One-to-one, haplotype-informed variant-calling against the Hg38 reference may ultimately be substituted with a pan-genome approach. Such methods are beginning to be implemented in a high-throughput fashion and are another means of representing population diversity (Ebler *et al*. 2022; Wang *et al*. 2023).

These high-resolution sequences will be of special value for the purposes of fine-mapping, which is especially relevant in the MHC, where long stretches of linkage disequilibrium have thwarted the uncovering of causal variants (Lam *et al*. 2015). Our release of 186 new ASW-like MHC haplotype assemblies will further assist in overcoming this challenge. This tool will also support the discovery of novel patterns of variation, and even never before-described genomic variation. Further, it is known that the unique sequence features of the MHC are driven by a combination of nuanced and interacting selective scenarios, converging on a single region of the genome (Hedrick 1998). With this new access to extended MHCs from large population cohorts, we can begin to more thoroughly reconstruct those processes, which, in turn, will lead to a better understanding of human disease. This method also has applications for non-human MHC sequences, which are of critical importance to conservation efforts of endangered species (Aguilar *et al*. 2004; Sommer 2005). The containerized, pipelined workflow established here will make its application to countless additional short-read datasets computationally accessible. In this way, MHConstructor will encourage functional interpretation of MHC variation and its roles in human disease contexts.

## Methods

### DNA collection for target-capture sequencing

DNA from healthy individuals (Oksenberg *et al*. 2004; Hollenbach *et al*. 2019) was obtained for target-capture sequencing. 183 healthy individuals from a population similar to the 1000GenomesProject Utah residents (CEPH) with Northern and Western European ancestry group and 299 healthy individuals from a population similar to the 1000GenomesProject African Ancestry in Southwest US group. Populations will be referred to as CEU-like and ASW-like, according to nomenclature recommendations from a National Academies consensus study report (National Academies of Sciences, Engineering, and Medicine; Division of Behavioral and Social Sciences and Education; Health and Medicine Division; Committee on Population; Board on Health Sciences Policy; Committee on the Use of Race, Ethnicity, and Ancestry as Population Descriptors in Genomics Research 2023)

### Target-capture sequencing of the extended MHC

For each sample, 100 ng of high-quality DNA was fragmented with the Library Preparation Fragmentation Kit 2.0 (Twist Bioscience, San Francisco, USA). Following fragmentation, DNA was end-repaired, and poly-A tails were attached and fragments then ligated to Illumina-compatible dual index adapters having unique barcodes. After ligating, fragments were purified with 0.8x ratio AMPure XP magnetic beads (Beckman Coulter, Brea, USA). Dual-size selection was performed (0.42x and 0.15x ratios), and libraries of approximately 800 bp were selected, amplified, and purified using magnetic beads.

Following fluorometric quantification, the samples were pooled (30 ng/sample) via ultrasonic acoustic energy. The Twist Target Enrichment kit (Twist Bioscience, San Francisco, USA) was then used to perform target capture on pooled samples. DNA libraries were bound to 33,620 biotinylated 120 bp probes designed to target the entire extended MHC, including genic and intergenic regions. Then, streptavidin magnetic beads were used to capture fragments targeted by the probes. Captured fragments were then amplified and purified. Bioanlyzer (Agilent, Santa Clara, USA) was then used to analyze the enriched libraries. After evaluation, enriched libraries were sequenced using a paired-end 150 bp sequencing protocol on the NovaSeq 6000 platform (Illumina, San Diego, USA). This protocol is an adaptation of the method previously published by Norman et al., 2017.

### WGS data collection from the 1000GenomesProject

The MHC region was extracted from 1000GenomesProject .cram files corresponding to 30x WGS data for 214 individuals from the CEU and ASW populations using the following command: $ samtools view -T http://ftp.1000genomes.ebi.ac.uk/vol1/ftp/technical/reference/GRCh38_reference_genome/GRCh38_full_analysis_set_plus_decoy_hla.fa -b -o <sample>.bam <sampleURL> <chromosome>, where <sample> and <sampleURL> correspond to the unique ID and location path on the ftp server for each individual and <chromosome> corresponds to the chromosome region from which to extract reads. Reads were extracted from the canonical MHC region using chr6: 28509120-33481577, as well as from the following alternate chromosomes on the hg38 build known to house MHC sequence: chr6_GL000250v2_alt, chr6_KI270800v1_alt, chr6_KI270799v1_alt, chr6_GL383533v1_alt, chr6_KI270801v1_alt, chr6_KI270802v1_alt, chr6_KB021644v2_alt, chr6_KI270797v1_alt, chr6_KI270798v1_alt, chr6_GL000251v2_alt, chr6_GL000252v2_alt, chr6_GL000253v2_alt, chr6_GL000254v2_alt, chr6_GL000255v2_alt, chr6_KI270758v1_alt. Reads which did not map to these MHC regions were extracted separately and used in later stages of analysis. The full list of 1000 Genomes Project individuals and the ftp links to access corresponding short-read fastq files can be found in Supplemental Document 1.

### Additional MHC-related tools

These processes are separate from the MHConstructor pipeline. They include two functions that produce data required for MHConstructor to run. However, these modules may be skipped if the user already has these data prepared.

- **HLA-DRB1 genotyping.** In order to assign each individual to the most relevant major MHC haplotype, according to Houwaart *et al*. 2023), we have included a pre-processing step to generate *HLA-DRB1* genotypes at first field resolution and complement component 4 (C4) genotype (C4A or C4B) and copy number. This stage is executed prior to the assembly algorithm. High throughput *HLA-DRB1* first field genotypes are generated using T1k (Song *et al*. 2023). Since *HLA-DRB1* is in known to be in strong linkage disequilibrium with *HLA-DRB3, -DRB4* and *-DRB5* (Rollini *et al*. 1985; Maiers *et al*. 2007), we infer that the first field genotypes of *HLA-DRB3, -DRB4* and *-DRB5* will correspond to previously established *HLA-DRB1* haplotypes (Houwaart *et al*. 2023). In this way, we determine the HLA class II haplotype(s) for each individual.
- **Complement component 4 (C4A and C4B genotyping.** *C4A* and *C4B* genotypes and copy numbers are assigned using C4Investigator (Marin *et al*. 2024) with the standard parameter values, calculating C4 copy numbers (C4A, C4B, C4S, C4L).

### MHConstructor *de novo* assembly pipeline

The following describes each functional module involved in MHConstructor. Each module is executed by a driver script written in bash, which calls the necessary software to execute the module. The modules associated with read quality filtering, contig generation and scaffolding are derived from the reference-guided *de novo* assembly method described by Lischer and Shimizu, 2017. For an in-depth description of these methods, please refer to the original publication (Lischer and Shimizu 2017). Changes made to the original algorithm include replacing the primary assembly software with *velvet* (Zerbino and Birney 2008) for increased speed, as well as minor code alterations, made to be applicable to human MHC sequence reads and up to date with current software versions.

**1. Short read quality control.** Fastq files are processed to remove stretches of homopolymer runs from reads, where reads contain a minimum of 20, consecutive guanines (G). Long, consecutive stretches of guanine is a known artifact of Illumina NGS (Stoler and Nekrutenko 2021). Then, reads are randomly sampled with a seed value using seqtk (https://github.com/lh3/seqtk) to select the desired number of reads to include in downstream analysis. Sampled reads are then processed with Trimmomatic (Bolger *et al*. 2014) to remove leading or trailing low quality bases (< 3 or N), remove bases that had an average quality score of 15 or less across four base pair sliding windows and exclude reads that are shorter than 40 base pairs, according to Lischer and Shimizu 2017. This analysis assumes that sequencing primers have already been removed from the sequencing reads.
**2. MHC BMH assignment and read binning.** Best matching haplotypes (BMHs) are inferred from HLA-DRB1 genotyping and C4 data. If an individual is homozygous at HLA-DRB1, they are assigned a single BMH. If an individual has a heterozygous DR-haplotype, they are assigned two BMHs. Possible recombination was not taken into account in assigning proxy chromosomal haplotypes. Once BMHs are assigned, reads are aligned to each BMH using *bowtie2* (Langmead and Salzberg 2012). The *bowtie2* minimum alignment score is set to “L,0,-0.6”, according to Step 2 of the reference-guided *de novo* assembly method in Lischer and Shimizu, 2017. Reads that align to a given BMH are then extracted for *de novo* assembly. Reads that do not align to the BMH are extracted and grouped for *de novo* assembly separate from the reads that aligned. If an individual has two BMHs, the *de novo* assembly process is conducted separately for each BMH. In these cases, following *de novo-*assembly, contigs generated from reads that did not map to a BMH are later aligned to the heterozygous individual’s alternative BMH and any cross-mapping contigs are removed, as described in section 3.
**3. Haplotype-informed contig set generation via reference-guided, de novo assembly.** Non-redundant supercontigs are generated using an individual’s BMH(s) as guide sequence(s) for *de novo* assembly, according to Lischer and Shimizu, 2017. Guide BMH sequences were obtained from the following NCBI accession numbers: OK649231.1, OK649232.1, OK649233.1, OK649234.1, OK649235.1, OK649236.1. and the MHC region on chromosome 6 of human reference genome GRCh38.p14, <u>NC_000006.12</u>: chr6: 28509120-33481577, Individuals with a DR8 haplotype were assigned GRCh38.p14. Processed reads are initially aligned to the guide BMH using *bowtie2* (Langmead and Salzberg 2012), with well-aligning reads being extracted and grouped into blocks and then overlapping superblocks. Reads that do not align to the guide BMH are extracted and assembled separately. Both categories of reads are then *de novo* assembled into contigs via de Bruijn kmer graphs, as implemented by *velvet* (Zerbino and Birney 2008). *Velvet* was chosen instead of other assemblers described in the original publication (Lischer and Shimizu 2017) for its use of de Bruijn graphs and because its faster speed makes it better suited to high throughput applications in the context of large disease cohorts. Reads not mapped to the reference sequence are *de novo* assembled separately. The resulting contigs from the un-mapped sequence are aligned to the rest of the human reference genome (HG38) and, for heterozygous individuals, to the alternative MHC haplotype, using *minimap2* (Li 2018). Any contigs that map outside of the MHC, or map to the alternative MHC haplotype reference are removed. This step is an addition to the original algorithm. The remaining filtered contigs generated from unmapped reads are then combined with contigs generated from mapped reads. Finally, overlapping contigs are then merged into non-redundant supercontigs, using *AMOS* (Treangen *et al*. 2011).
**4. Error-correcting, *de novo* scaffolding, and gap closing of haplotype-informed assemblies** Original reads are mapped onto the contigs and *GATK HaplotypeCaller* (McKenna *et al*. 2010; DePristo *et al*. 2011; Van der Auwera *et al*. 2013) is used to perform localized error-correction. This represents a deviation from the tool described in the original publication (Lischer and Shimizu 2017), as the GATK error-correction tool originally chosen is no longer supported by GATK. Corrected contigs are then aligned to BMH using *nucmer* and the resulting delta output files are used as input to *Assemblytics* (Nattestad and Schatz 2016), which establishes contig orientation and identified structural variants. Haplotype-informed contig sets are then processed into draft assemblies, via the Lischer and Shimizu algorithm (Lischer and Shimizu 2017). *SOAPdenovo2* (Luo *et al*. 2012) is used to scaffold error-corrected contigs and close gaps using paired-end mapping to create the haplotype-informed assembly.
**5. Haplotype-informed assembly completion and scaffold orienting.** Draft scaffolds are then ordered and oriented with respect to their major MHC phased reference sequence (Houwaart *et al*. 2023) using RagTag (Alonge *et al*. 2022). A final assembly fasta file is created, which contains one single fasta record representing the ordered and continuous sequence assembled with respect to the reference, as well as individual fasta records representing the scaffolds that RagTag did not place on MHC guide haplotype sequence. This serves as a haplotype-informed consensus sequence for each major MHC haplotype.

### Post-hoc assembly parameter evaluation and optimization

Following algorithm development, the effect of read count on assembly time, computational cost, and assembly quality was evaluated by randomly subsampling pairs of reads at 200k, 500k, 1M, 2M, and 3M intervals for each sample. Additionally, to evaluate the most appropriate kmer length for *de novo* assembly, *Kmergenie* (Chikhi and Medvedev 2014) was used to estimate initial ranges of likely successful k-mers. This was done separately for the subset of mapped reads and the subset of reads that did not align to the reference. Paired R1 and R2 reads were combined for these estimates. Best kmer predictions were averaged across all samples at a given read count. The average kmer size prediction was then rounded to the closest whole, odd number. Standard deviation of best predicted kmer was also evaluated across all samples. Time, memory usage and quality were measured separately for super contig and scaffolding stages.

### Assembly method validation using high-quality MHC reference sequences

To validate the accuracy of assemblies generated with this method, we applied it to independently generated Illumina short-read datasets, corresponding to the MHC region of HLA-homozygous, consanguineous cell lines (*SRA:* SRP348947, *BioProject:* <u>PRJNA764575</u>) (Norman *et al*. 2017). We chose five cell lines that were also thoroughly assembled and curated in a later study via a combination approach using both short and long-read sequencing technology (Houwaart *et al*. 2023). data from individuals from the30xthat had their MHC phased sequence assembled with - technology Following assembly of these five short read datasets using our own *de novo* approach, we aligned our new assemblies to the curated, full-length MHC phased sequence for that same cell line, acting as the gold standard. Any variation with respect to the reference sequence was counted as an error. Base pair accuracy validation was performed using *minimap2* to align MHConstructor sequences to the corresponding reference sequence and calling SNPs using *paftools*. Structural errors >=1bp INDELs were identified using the assembly-based variant predictor *Assemblytics* (Nattestad and Schatz 2016). The nucleotide length of each error was summed for each size category and a final percent error was calculated for each assembly by dividing the number of erroneous base pairs in each category by the full length of assembled sequence. To evaluate the impact of heterozygous, diploid sequence data on assembly accuracy, we used three 1000GenomesProject 30X WGS samples with long read data that has recently been assembled into phased MHC haplotypes (Huijse *et al*. 2023).

### Repeat annotation and masking

To characterize the repetitive elements located within novel MHC assemblies, we applied repeatMasker v4 (http://www.repeatmasker.org) to the consensus sequence(s) from each assembly, using default parameters. Following characterization, repeat regions were then masked.

## Declarations

### Ethics approval and consent to participate

Sequencing data was granted exemption due to deidentified status.

### Consent for publication

Not applicable.

### Availability of data and materials

MHConstructor is publicly available at www.github.com/Hollenbach-lab/MHConstructor. Data for this project is available at the NCBI record phs003244. The BioProject for this record includes links to SRA, short read fastq data, *de novo* assembly fasta and scaffolded, consensus sequence fasta, per individual. Additional intermediate data generated throughout the *de novo* assembly process is available upon request.

### Authors’ contributions

Study conceptualization: KW, JO, PN, DA, JH; novel software development: KW, RS; software implementation: KW, RS, AR, AS; data generation: KW, RS, KK, JW, JB, SC, DA; data analysis: KW, RS, KK, JW, JB; data interpretation: KW, RS, NP, PN, DA, JH; manuscript writing: KW, RS, JH; manuscript revision: KW, RS, JW, NP, AS, JO, PN, DA, JH.

## Acknowledgments

DNA samples were provided by the UCSF MS Biorepository supported by grant SI-2001-35701(JO) from the National Multiple Sclerosis Society. We are very grateful for the contributions of Sully Adams, Julianne David, Lisa Huijse and Solomon Endlich from Base5 Genomics, Inc in providing us access to their pre-print data, as well as for helpful discussion. We acknowledge Ting Wang, David Sayer, Stephan Scholz and Alexander Dilthey for their valuable commentary and discussions during the development of this method. We would also like to thank Tony Capra for his software suggestion.

