## Supplemental Material for "MHConstructor: A high-throughput, haplotype-informed solution to the MHC assembly challenge"

### Supplemental data

#### Table of contents:


#### Results

##### 1. Haplotype-guided, *de novo* short read MHC assembly workflow

##### 2. Evaluation of assembly performance metrics

I. **Supplemental Table 1.** Best kmer size predictions for mappable reads, at given read counts.

II. **Supplemental Table 2.** Best kmer size predictions for un-mappable reads, at given read counts.

III. **Supplemental Figure S1.** Impact of kmer size choice on *de novo* assembly contig metrics, for mapped read k= 41, k=51 and k=61.

IV. **Supplemental Table S3.** User hours to generate assemblies. mappable read k = 49, unmappable read k=21.

V. **Supplemental Table S4.** User time (in minutes) to generate contigs and scaffold assemblies in target-capture short reads across multiple coverage depths.

VI. **Supplemental Figure S2.** Impact of starting read count on *de novo* assembly contig metrics.

##### 3. Quantification of MHConstructor error rates using high-quality MHC reference sequences

VII. **Supplemental Table S5.** Assembly statistics from Norman *et al.*, 2017 MHC reference haplotype Illumina short read dataset. *SRA:* SRP348947, *BioProject:* PRJNA764575.

VIII. **Supplemental table S6.** Reference MHC coverage with contigs generated from reads that mapped to reference genome.

IX. **Supplemental Table S7.** Reference MHC coverage with contigs generated from reads that did not map to initial reference genome.

X. **Supplemental Figure S3.** Dotplot of *Assemblytics* assembly alignments between 1KGP WGS MHConstructor assemblies (y-axes) and their corresponding, phased reference MHC haplotypes (x-axes).

XI. **Supplemental Figure S4.** *Assemblytics* analysis of false SV composition for the MHConstructor *de novo* assembly of HG00621, as compared to phased, reference MHC haplotypes for HG00621.

XII. **Supplemental Figure S5.** *Assemblytics* analysis of false SV composition for the MHConstructor *de novo* assembly of NA19240, as compared to phased, reference MHC haplotypes for NA19240.

XIII. **Supplemental Figure S6.** *Assemblytics* analysis of false SV composition for the MHConstructor *de novo* assembly of HG00621, as compared to phased, reference MHC haplotypes for NA20129.

XIV. **Supplemental Table S8**. Number of error sites in base5-validated *de novo* assemblies that are caused by inclusion of a heterozygous site belonging to the alternative haplotype and those caused by assembly error.

XV. **Supplemental figure S7.** Mauve alignment plots confirm that the two DR haplotypes for heterozygous Individual Z are correctly assembled.

XVI. **Supplemental Figure S8.** HLA Class II structure can be correctly assembled even when an incorrectly-match haplotype is used as the assembly guide, so long as the correct haplotype is used for scaffolding orientation

#### 4. MHConstructor assembly of target-capture 60x and 1000genomes 30x WGS short read MHC data.

XVII. **Supplemental Figure S9**. MHConstructor assembly metrics for target capture and WGS assemblies.

XVIII. **Supplemental Figure S10.** Nucleotide diversity within sequencing reads.

**Supplemental Table 1.** Best kmer size predictions for mappable reads, at given read counts.

| ID | 200k | 500k | 1M | 2M | 3M |
|---|---|---|---|---|---|
| EPIC0046 | 17 | No best k | 31 | 45 | 51 |
| EPIC0053 | 19 | No best k | 25 | 61 | 33 |
| EPIC0176 | 17 | No best k | 21 | 35 | 71 |
| EPIC0242 | 19 | No best k | 51 | 45 | 71 |
| EPIC0246 | 27 | 21 | 21 | 57 | 25 |
| Average | 19.8 | NA | 29.8 | 48.6 | 50.2 |
| StDev | 4.15 | NA | 12.54 | 10.43 | 21.19 |

**Supplemental Table 2.** Best kmer size predictions for un-mappable reads, at given read counts.

| ID | 200k | 500k | 1M | 2M | 3M |
|---|---|---|---|---|---|
| EPIC0046 | 17 | 33 | 17 | 31 | 31 |
| EPIC0053 | 31 | 23 | 23 | 31 | 17 |
| EPIC0176 | 17 | 17 | 23 | 19 | 19 |
| EPIC0242 | 27 | 19 | 23 | 21 | 23 |
| EPIC0246 | 36 | 23 | 27 | 21 | 17 |
| Average | 25.6 | 23 | 22.6 | 24.6 | 21.4 |
| StDev | 8.47 | 6.16 | 3.58 | 5.90 | 5.90 |

A.

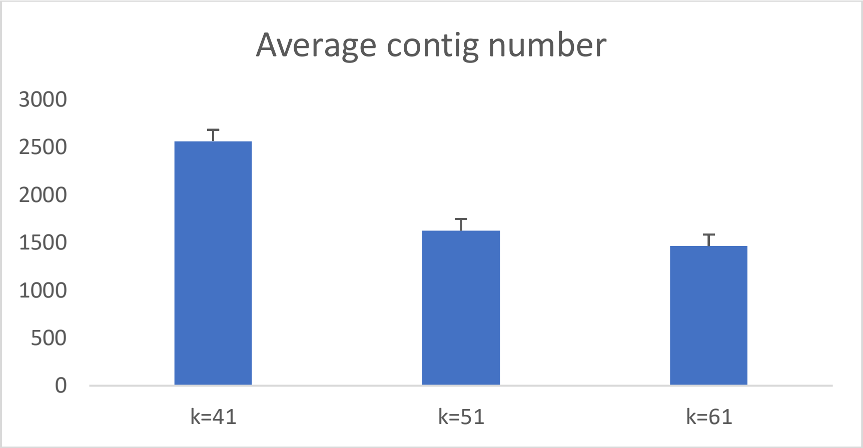


B.

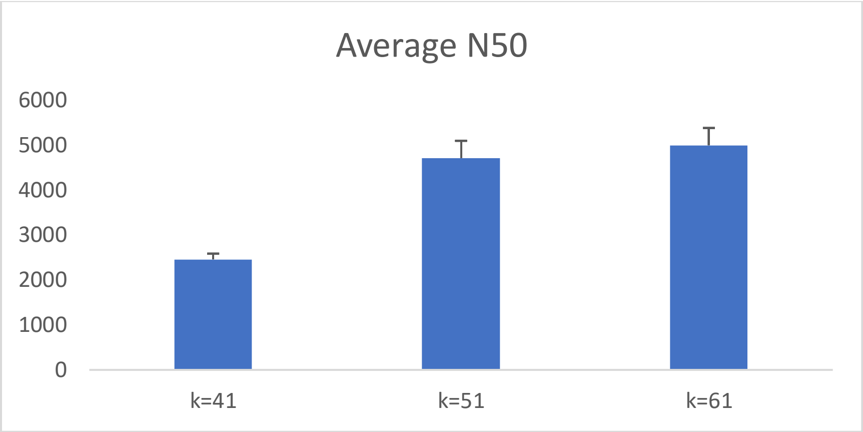


C.

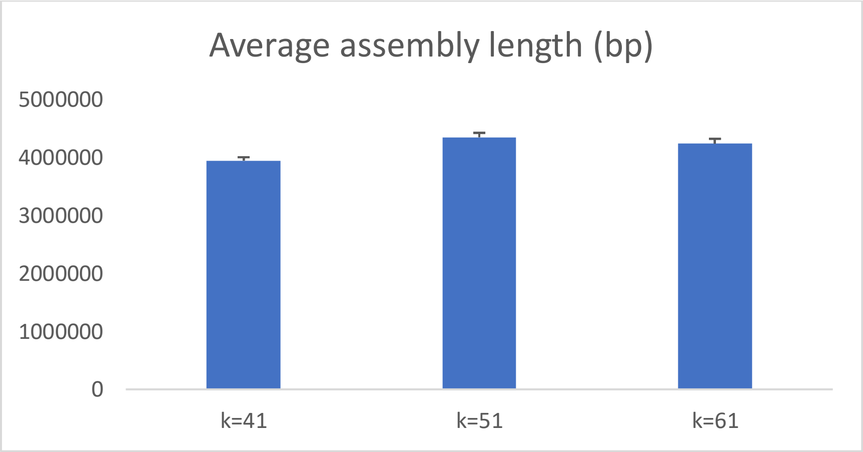


**Supplemental Figure S1.** Impact of kmer size choice on *de novo* assembly contig metrics, for mapped read k= 41, k=51 and k=61. Test samples subsampled to contain 2 million reads per individual, n=14. A) Average number of contigs generated for each set of subsampled reads. B) Average N50 length (in bp) for each set of subsampled reads. C) Average length (bp) of contigs generated by *de novo* assembly.

**Supplemental Table S3.** User hours to generate assemblies. mappable read k = 49, unmappable read k=21.

| Sample ID | 200k | 500k | 1M | 2M | 3M |
|---|---|---|---|---|---|
| EPIC0046 | 0.07 | 0.34 | 0.89 | 2.36 | 6.21 |
| EPIC0053 | 0.09 | 0.21 | 0.66 | 1.68 | 3.99 |
| EPIC0176 | 0.07 | 0.21 | 0.65 | 1.84 | 6.07 |
| EPIC0242 | 0.08 | 0.24 | 0.55 | 2.16 | 2.85 |
| EPIC0246 | 0.09 | 0.23 | 0.57 | 1.63 | 4.85 |
| Average | 0.08 | 0.25 | 0.66 | 1.93 | 4.79 |

**Supplemental Table S4.** User time (in minutes) to generate contigs and scaffold assemblies in target-capture short reads across multiple coverage depths.

| Average coverage | Starting reads | Assembly | Scaffolding+EC | Total |
|---|---|---|---|---|
| ~25X | 2 million | 36 | 24 | 61 |
| ~45X | 3 million | 41 | 40 | 81 |
| ~60X | 4 million | 53 | 51 | 104 |

Performed on HPC cluster, 8 threads per sample

A.

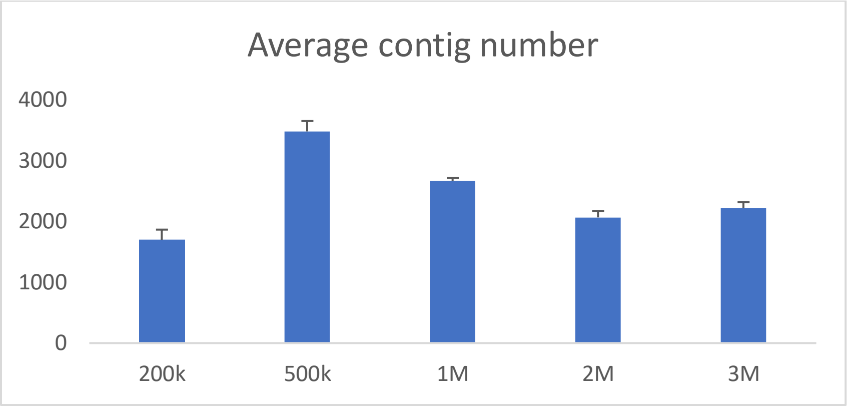


B.

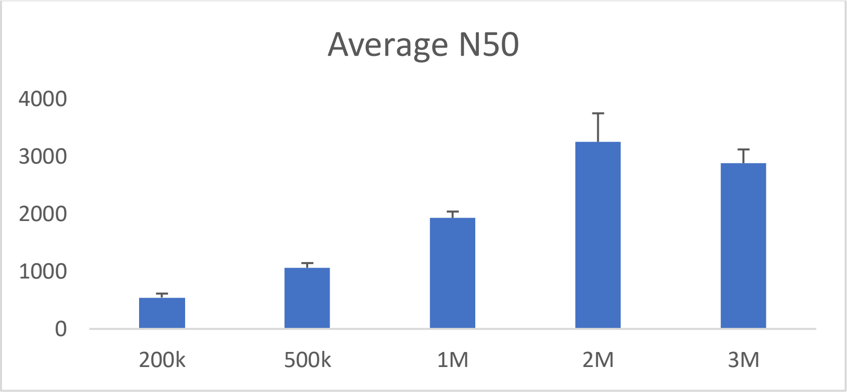


C.

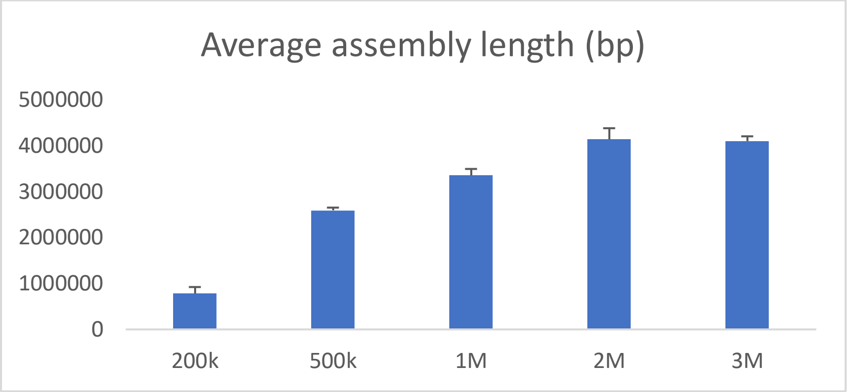


**Supplemental Figure S2.** Impact of starting read count on *De novo* assembly contig metrics. Metrics were generated for a test dataset (n=5) of randomly sampled, starting read counts: 200,000 (200k), 500,000 (500k), 1 million (1M), 2 million (2M) and 3 million (3M). A) Average number of contigs generated for each set of subsampled reads. B) Average N50 length (in bp) for each set of subsampled reads. C) Average length (bp) of contigs generated by *de novo* assembly.

**Supplemental Table S5.** Assembly statistics from Norman *et al.*, 2017 MHC reference haplotype Illumina short read dataset. *SRA:* SRP348947, *BioProject:* PRJNA764575.

| Assembly metrics | OK649231 | OK649232 | OK649234 | OK649235 | OK649236 |
|---|---|---|---|---|---|
| Number of reads | 266995 | 217603 | 304342 | 210039 | 252657 |
| Contigs (n) | 1506 | 2227 | 1147 | 2425 | 1585 |
| N50 | 4055 | 2549 | 6191 | 2274 | 3964 |
| Max contig (bp) | 24727 | 18382 | 36889 | 16305 | 23271 |
| Total assembly length (bp) | 4127329 | 3908195 | 4539318 | 3710459 | 4196340 |


| Assembly metrics | OK649231 | OK649232 | OK649234 | OK649235 | OK649236 |
|---|---|---|---|---|---|
| Number of reads | 266995 | 217603 | 304342 | 210039 | 252657 |
| Contigs (n) | 1506 | 2227 | 1147 | 2425 | 1585 |
| N50 | 4055 | 2549 | 6191 | 2274 | 3964 |
| Max contig (bp) | 24727 | 18382 | 36889 | 16305 | 23271 |
| Total assembly length (bp) | 4127329 | 3908195 | 4539318 | 3710459 | 4196340 |

**Supplemental table S6.** Reference MHC coverage with contigs generated from reads that mapped to reference genome

| MHC Haplotype | % of haplotype covered | Total haplotype covered (bp) | Average % Identity | Median % Identity |
|---|---|---|---|---|
| APD | 72 | 3591688 | 99.6 | 100 |
| DBB | 59.48 | 3002824 | 99.7 | 100 |
| MANN | 80 | 4022500 | 99.53 | 100 |
| QBL | 60.51 | 2968060 | 99.37 | 100 |

**Supplemental Table S7.** Reference MHC coverage with contigs generated from reads that did not map to initial reference genome.

| MHC Haplotype | % Haplotype covered | Total haplotype coverage (bp) | Average % Identity | Median % Identity |
|---|---|---|---|---|
| APD | 0.078 | 3872 | 99.8 | 100 |
| DBB | 0.140 | 7153 | 99.8 | 100 |
| MANN | 0.160 | 8233 | 99.83 | 100 |
| QBL | 0.053 | 2611 | 99.54 | 100 |

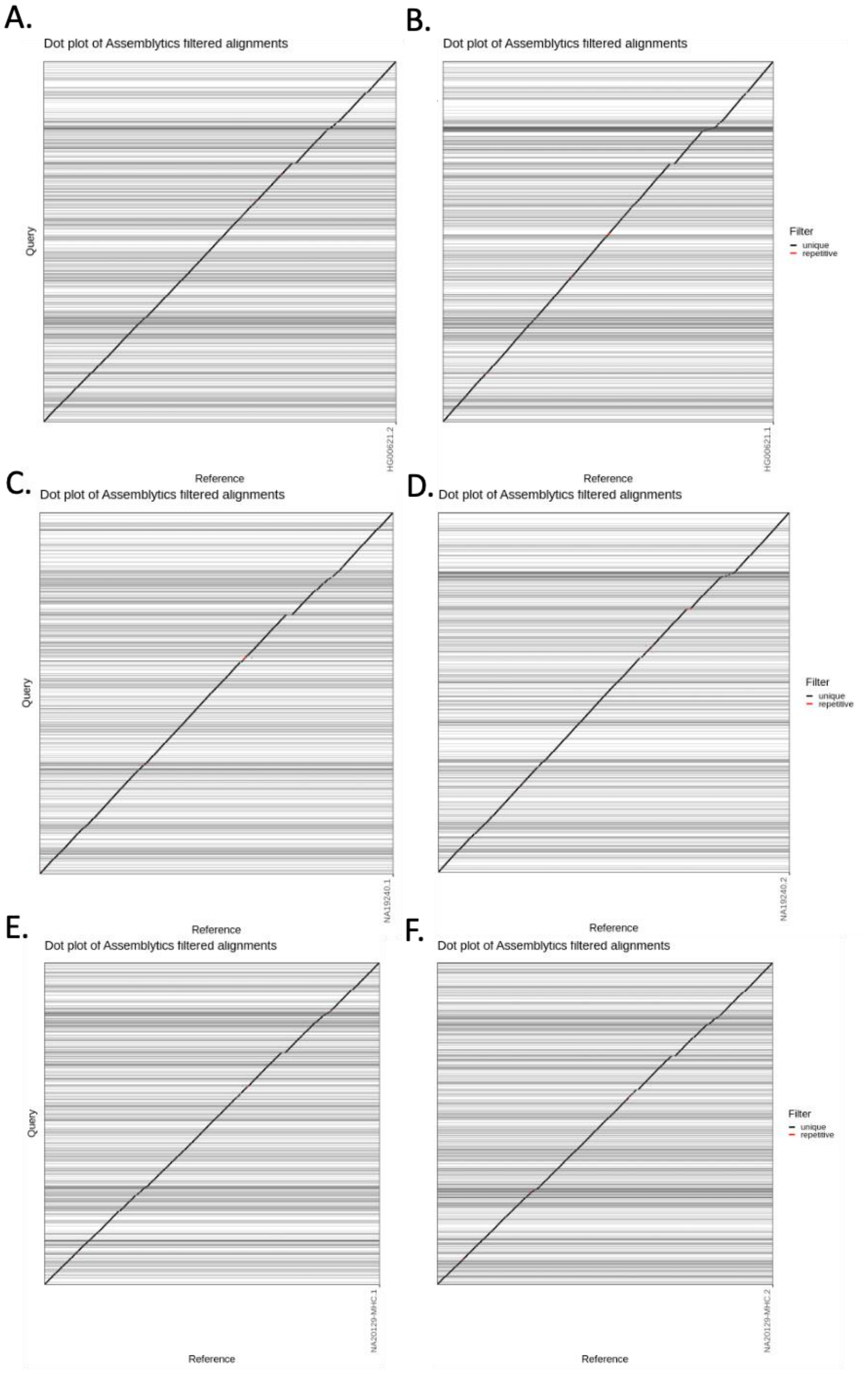


**Supplemental Figure S3.** Dotplot of *Assemblytics* assembly alignments between 1KGP WGS MHConstructor assemblies (y-axes) and their corresponding, phased reference MHC haplotypes (x-axes). The assemblies shown represent: HG00621 MHC haplotype A (A), HG00621 MHC haplotype B (B), NA19240 MHC haplotype A (C), NA19240 MHC haplotype B (D), NA20129 MHC haplotype A (E), NA20129 MHC haplotype B (F).

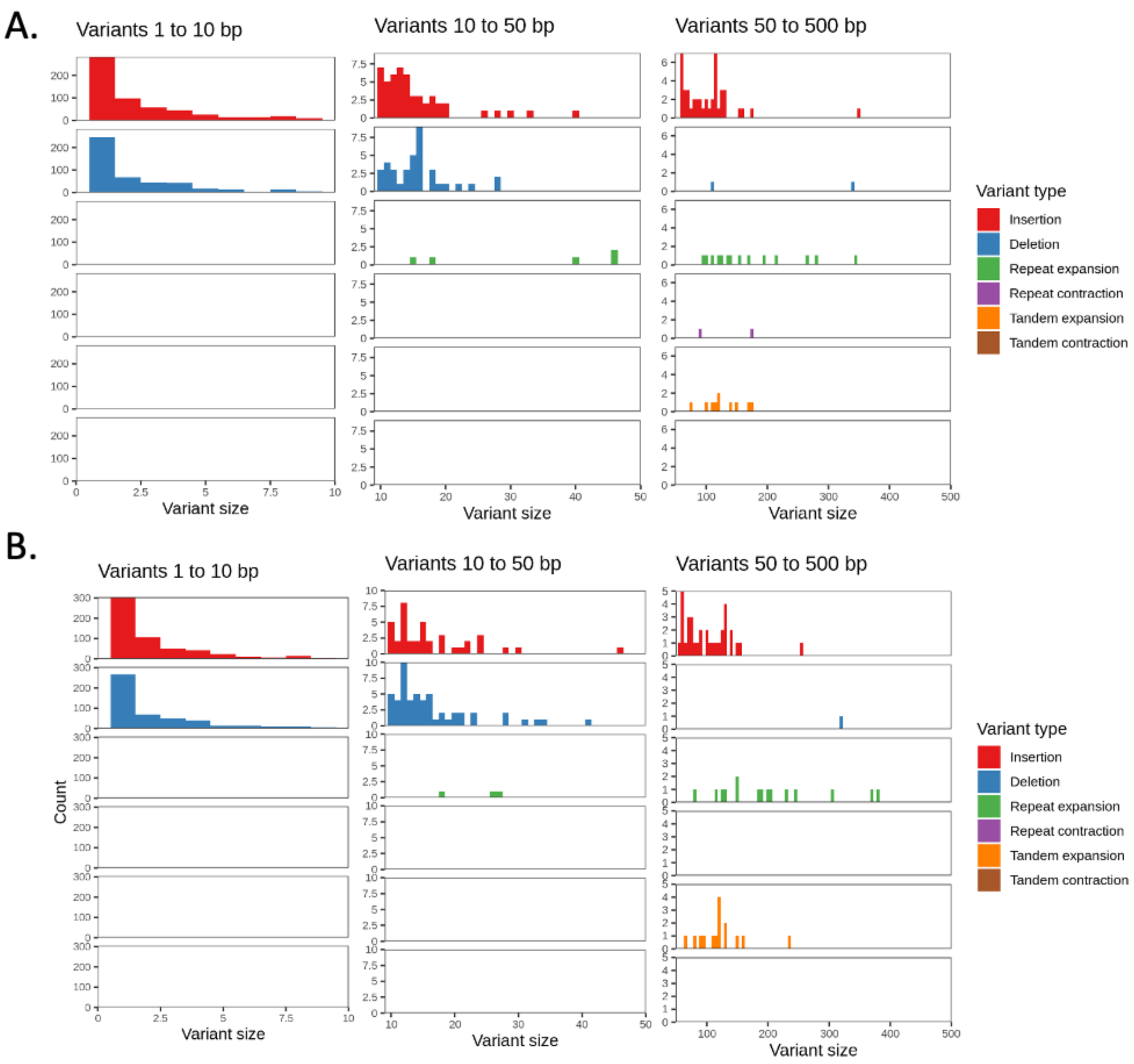


**Supplemental Figure S4.** *Assemblytics* analysis of false SV composition for the MHConstructor *de novo* assembly of HG00621, as compared to phased, reference MHC haplotypes for HG00621. A) HG00621 MHC haplotype A, B) HG00621 MHC haplotype B. Categories of assembly SV errors described in legend.

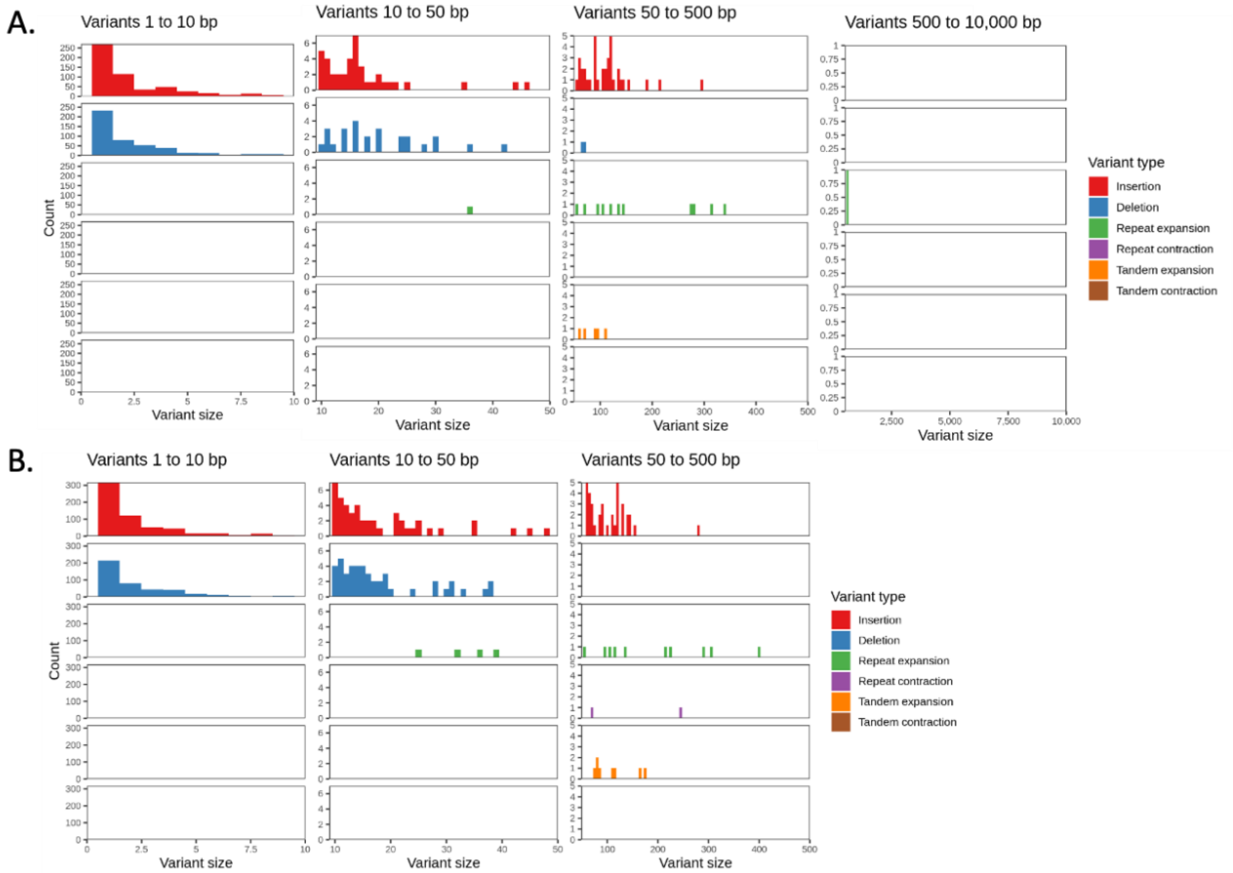


**Supplemental Figure S5.** *Assemblytics* analysis of false SV composition for the MHConstructor *de novo* assembly of NA19240, as compared to phased, reference MHC haplotypes for NA19240. A) NA19240 MHC haplotype A, B) NA19240 MHC haplotype B. Categories of assembly SV errors described in legend.

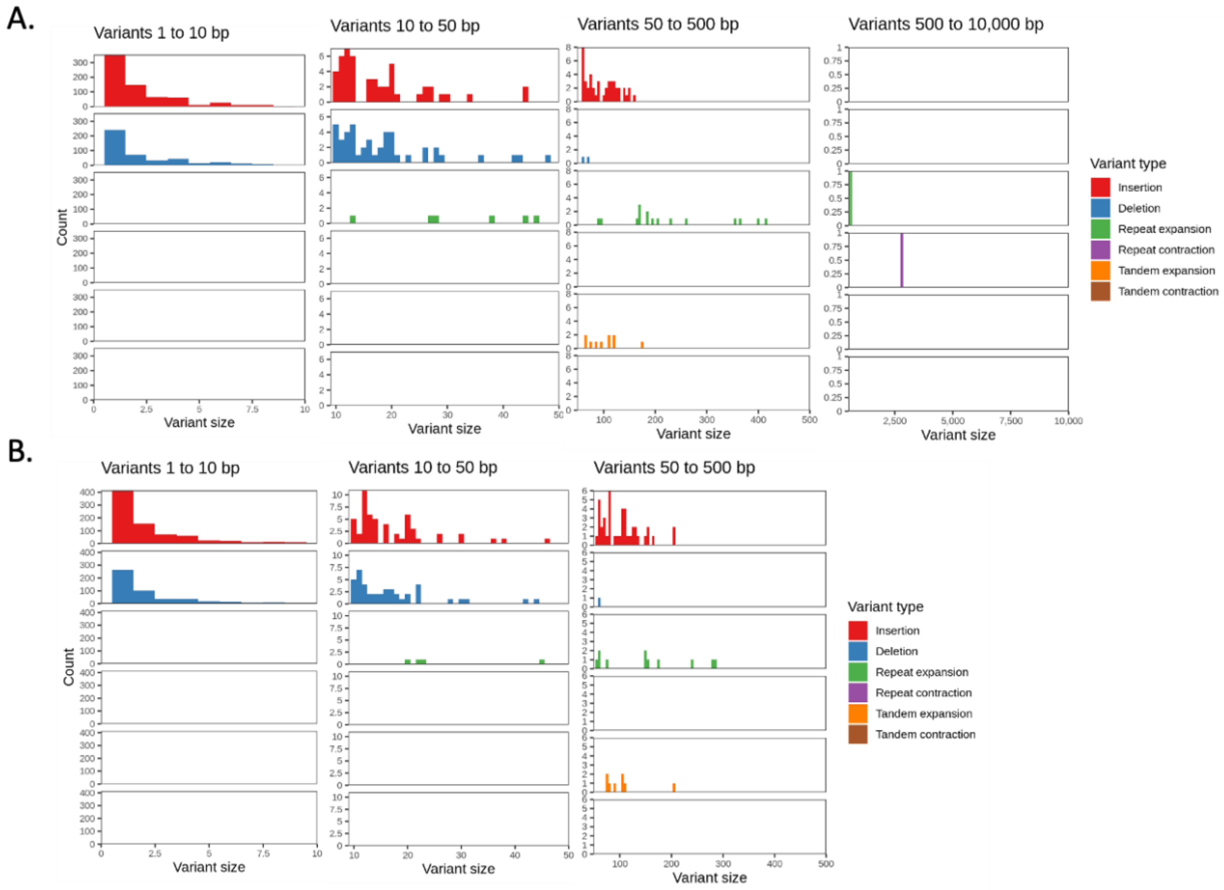


**Supplemental Figure S6.** *Assemblytics* analysis of false SV composition for the MHConstructor *de novo* assembly of HG00621, as compared to phased, reference MHC haplotypes for NA20129. A) NA20129 MHC haplotype A, B) NA20129 MHC haplotype B. Categories of assembly SV errors described in legend.

**Supplemental Table S8**. Number of error sites in base5-validated *de novo* assemblies that are caused by inclusion of a heterozygous site belonging to the alternative haplotype and those caused by assembly error.

| Site category | HG00621.1 | NA19240.1 | NA20129.1 |
|---|---|---|---|
| Heterozygous sites | 5167 | 4602 | 5728 |
| Assembly error sites | 3068 | 5447 | 3021 |

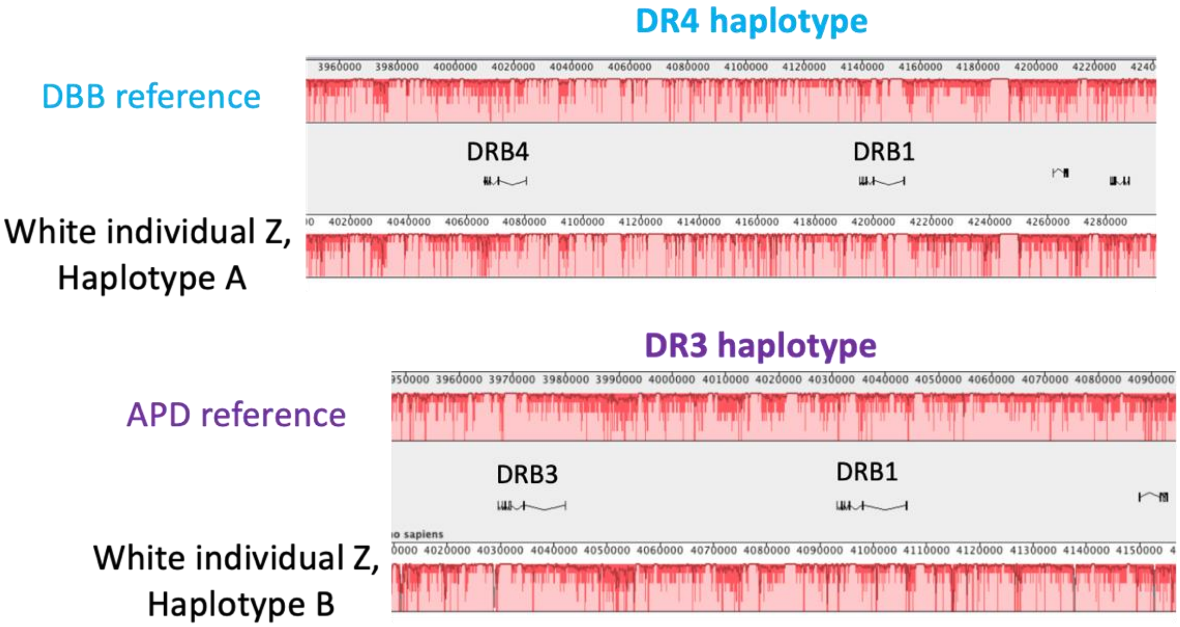


**Supplemental figure S7.** Mauve alignment plots confirm that the two DR haplotypes for heterozygous Individual Z are correctly assembled. In the top plot, the Individual Z sequence corresponding to the DR4 haplotype is displayed aligned against the DBB (DR4) published reference sequence (Houwaart *et al.*, 2022). In the bottom plot, the Individual Z sequence corresponding to the DR3 haplotype is displayed aligned against the APD (DR3) published reference sequence (Houwaart *et al.*, 2022). Light red indicates perfect alignment, dark red indicates presence of substitutions. HLA-DRB genes are labelled above their annotated coding regions.

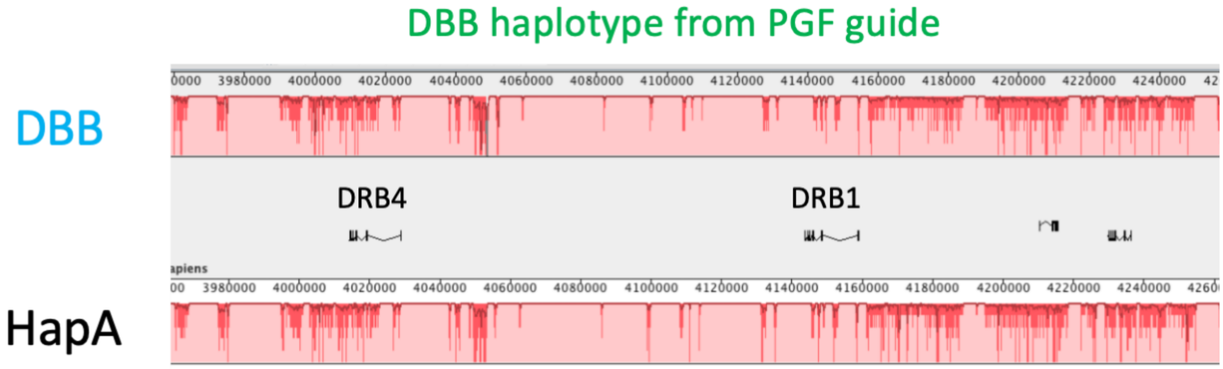


**Supplemental Figure S8.** HLA Class II structure can be correctly assembled even when an incorrectly-match haplotype is used as the assembly guide, so long as the correct haplotype is used for scaffolding orientation.

**Supplemental Table S9.** RepeatMasker annotation of repetitive sequence content in an MHConstructor WGS assembly and its corresponding phased MHC haplotype.

##### MHConductor NA19240.1

GC level: 44.77 %
bases masked: 2744988 bp ( 53.95 %)

| | number of elements* | length occupied | percentage of sequence |
|---|---|---|---|
| SINEs: | 3396 | 821766 bp | 16.15 % |
| ALUs | 2798 | 728111 bp | 14.31 % |
| MIRs | 592 | 92718 bp | 1.82 % |
| LINEs: | 1731 | 1050336 bp | 20.64 % |
| LINE1 | 1134 | 848383 bp | 16.67 % |
| LINE2 | 524 | 181556 bp | 3.57 % |
| L3/CR1 | 51 | 12950 bp | 0.25 % |
| LTR elements: | 992 | 617160 bp | 12.13 % |
| ERVL | 228 | 171551 bp | 3.37 % |
| ERVL-MaLRs | 358 | 152149 bp | 2.99 % |
| ERV_classI | 295 | 215025 bp | 4.23 % |
| ERV_classII | 47 | 56590 bp | 1.11 % |
| DNA elements: | 700 | 174889 bp | 3.44 % |
| hAT-Charlie | 396 | 93293 bp | 1.83 % |
| TcMar-Tigger | 151 | 54356 bp | 1.07 % |
| Unclassified: | 21 | 13236 bp | 0.26 % |
| Total interspersed repeats: | | 2677387 bp | 52.62 % |
| Small RNA: | 87 | 7042 bp | 0.14 % |
| Satellites: | 4 | 3266 bp | 0.06 % |
| Simple repeats: | 991 | 45653 bp | 0.90 % |
| Low complexity: | 191 | 11161 bp | 0.22 % |

##### Base5 NA19240.1

GC level: 44.90 %
bases masked: 2895112 bp ( 57.11 %)

| | number of elements* | length occupied | percentage of sequence |
|---|---|---|---|
| SINEs: | 3446 | 849271 bp | 16.75 % |
| ALUs | 2828 | 751675 bp | 14.83 % |
| MIRs | 611 | 96462 bp | 1.90 % |
| LINEs: | 1761 | 1122194 bp | 22.14 % |
| LINE1 | 1138 | 906985 bp | 17.89 % |
| LINE2 | 545 | 193494 bp | 3.82 % |
| L3/CR1 | 57 | 14535 bp | 0.29 % |
| LTR elements: | 1019 | 649642 bp | 12.81 % |
| ERVL | 232 | 177122 bp | 3.49 % |
| ERVL-MaLRs | 370 | 160049 bp | 3.16 % |
| ERV_classI | 299 | 228338 bp | 4.50 % |
| ERV_classII | 48 | 61061 bp | 1.20 % |
| DNA elements: | 728 | 182701 bp | 3.60 % |
| hAT-Charlie | 413 | 97084 bp | 1.91 % |
| TcMar-Tigger | 149 | 56228 bp | 1.11 % |
| Unclassified: | 20 | 21501 bp | 0.42 % |
| Total interspersed repeats: | | 2825309 bp | 55.73 % |
| Small RNA: | 90 | 7288 bp | 0.14 % |
| Satellites: | 7 | 3455 bp | 0.07 % |
| Simple repeats: | 1013 | 46435 bp | 0.92 % |
| Low complexity: | 202 | 12146 bp | 0.24 % |

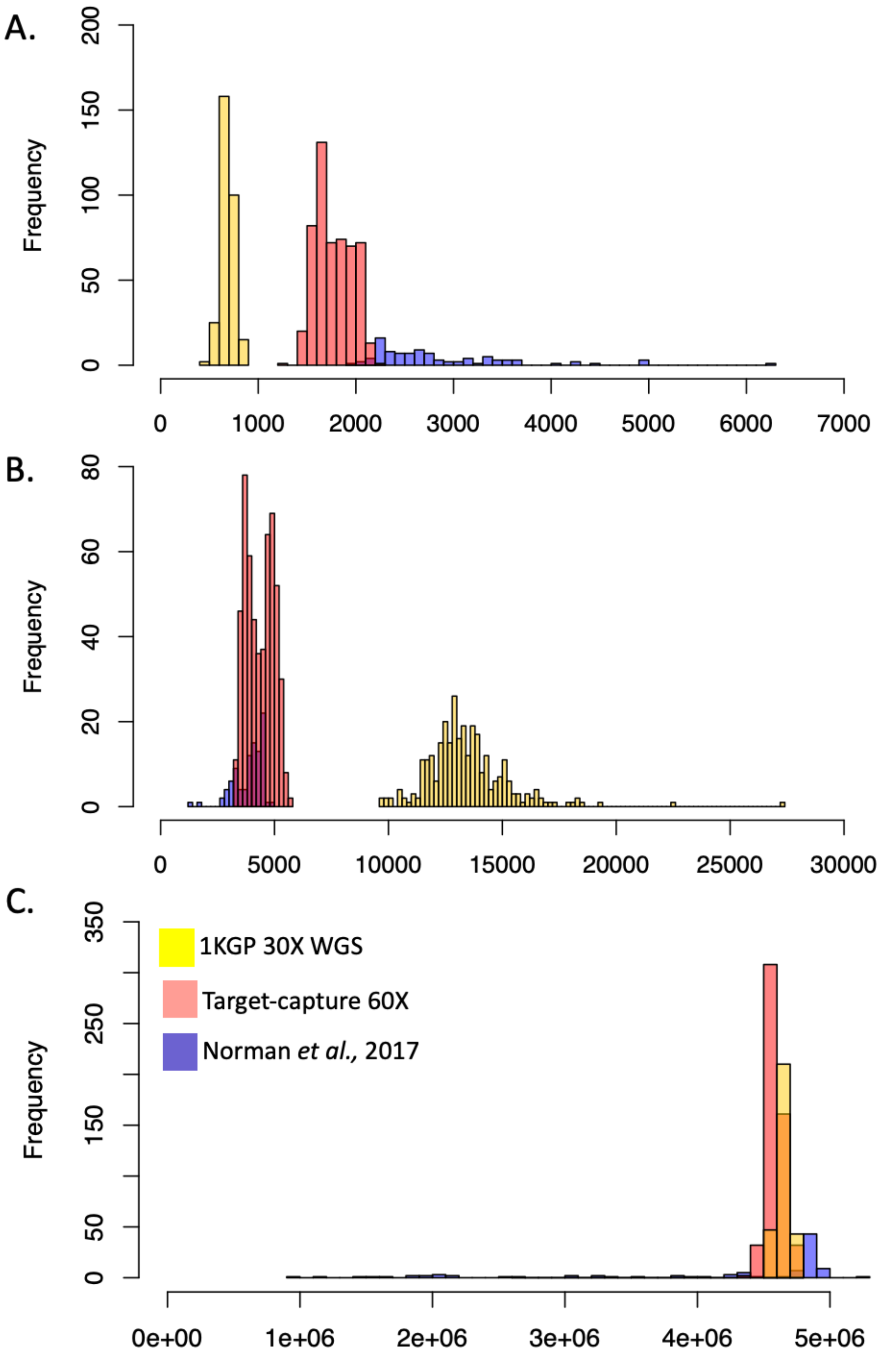


**Supplemental Figure S9. MHConstructor assembly metrics for target capture and WGS assemblies.** Purple distributions represent published results from Norman *et al.*, 2017, n=95 haplotypes. Pink distributions represent assemblies generated from target-capture, 60x short read data, n=536 haplotypes. Yellow distributions represent assemblies generated from 1000Genomes WGS 30x short read data, n=300 haplotypes. A) Histogram of the number of contigs per assembly. B) Histogram of assembly N50 (bp), per assembly. C) Histogram of assembly lengths (bp).

.

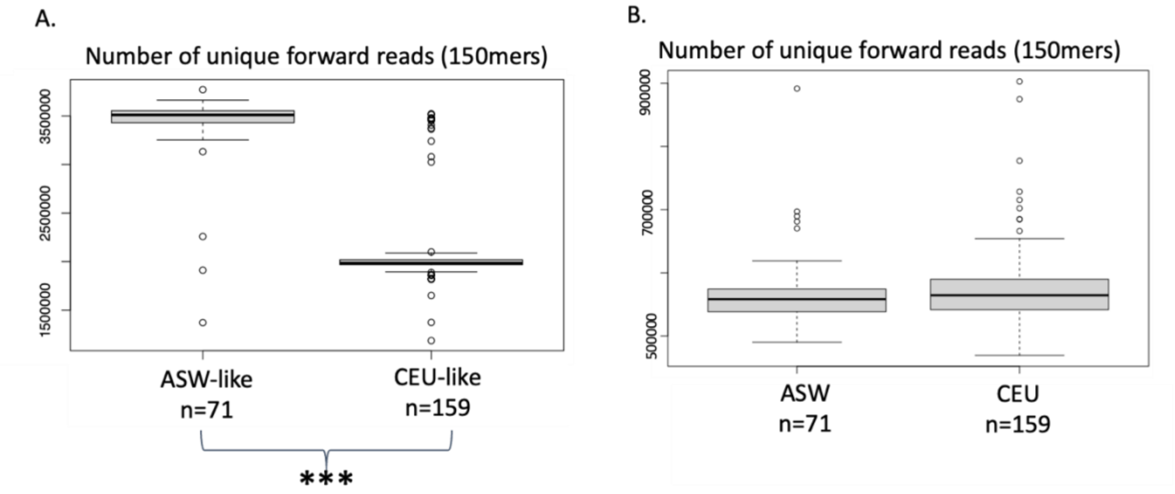


**Supplemental Figure S10.** Nucleotide diversity within sequencing reads. A) Number of unique forward reads in subsampled ASW-like and CEU-like individuals, from target-capture approach. B) Number of unique forward reads in ASW and CEU populations, from WGS approach.
