## Supplemental Document 1 for "MHConstructor: A high-throughput, haplotype-informed solution to the MHC assembly challenge"

**Individuals with 30X WGS data in the 1000GenomeProject African Ancestry in Southwest US group (ASW)**

ftp://ftp.sra.ebi.ac.uk/vol1/run/ERR323/ERR3239893/NA19625.final.cram

ftp://ftp.sra.ebi.ac.uk/vol1/run/ERR323/ERR3239917/NA19707.final.cram

ftp://ftp.sra.ebi.ac.uk/vol1/run/ERR323/ERR3239963/NA19900.final.cram

ftp://ftp.sra.ebi.ac.uk/vol1/run/ERR323/ERR3239966/NA19908.final.cram

ftp://ftp.sra.ebi.ac.uk/vol1/run/ERR323/ERR3239971/NA19920.final.cram

ftp://ftp.sra.ebi.ac.uk/vol1/run/ERR323/ERR3239973/NA19982.final.cram

ftp://ftp.sra.ebi.ac.uk/vol1/run/ERR323/ERR3239978/NA20281.final.cram

ftp://ftp.sra.ebi.ac.uk/vol1/run/ERR323/ERR3239985/NA20299.final.cram

ftp://ftp.sra.ebi.ac.uk/vol1/run/ERR323/ERR3239992/NA20346.final.cram

ftp://ftp.sra.ebi.ac.uk/vol1/run/ERR324/ERR3240095/NA19922.final.cram

ftp://ftp.sra.ebi.ac.uk/vol1/run/ERR324/ERR3240101/NA20412.final.cram

ftp://ftp.sra.ebi.ac.uk/vol1/run/ERR323/ERR3239914/NA19701.final.cram

ftp://ftp.sra.ebi.ac.uk/vol1/run/ERR323/ERR3239919/NA19712.final.cram

ftp://ftp.sra.ebi.ac.uk/vol1/run/ERR323/ERR3239920/NA19713.final.cram

ftp://ftp.sra.ebi.ac.uk/vol1/run/ERR323/ERR3239968/NA19914.final.cram

ftp://ftp.sra.ebi.ac.uk/vol1/run/ERR323/ERR3239975/NA20127.final.cram

ftp://ftp.sra.ebi.ac.uk/vol1/run/ERR323/ERR3239980/NA20287.final.cram

ftp://ftp.sra.ebi.ac.uk/vol1/run/ERR323/ERR3239987/NA20317.final.cram

ftp://ftp.sra.ebi.ac.uk/vol1/run/ERR323/ERR3239989/NA20334.final.cram

ftp://ftp.sra.ebi.ac.uk/vol1/run/ERR323/ERR3239994/NA20356.final.cram

ftp://ftp.sra.ebi.ac.uk/vol1/run/ERR323/ERR3239913/NA19700.final.cram

ftp://ftp.sra.ebi.ac.uk/vol1/run/ERR323/ERR3239918/NA19711.final.cram

ftp://ftp.sra.ebi.ac.uk/vol1/run/ERR323/ERR3239967/NA19909.final.cram

ftp://ftp.sra.ebi.ac.uk/vol1/run/ERR323/ERR3239972/NA19921.final.cram

ftp://ftp.sra.ebi.ac.uk/vol1/run/ERR323/ERR3239974/NA20126.final.cram

ftp://ftp.sra.ebi.ac.uk/vol1/run/ERR324/ERR3240096/NA19923.final.cram

ftp://ftp.sra.ebi.ac.uk/vol1/run/ERR323/ERR3239979/NA20282.final.cram

ftp://ftp.sra.ebi.ac.uk/vol1/run/ERR323/ERR3239986/NA20314.final.cram

ftp://ftp.sra.ebi.ac.uk/vol1/run/ERR323/ERR3239993/NA20348.final.cram

ftp://ftp.sra.ebi.ac.uk/vol1/run/ERR324/ERR3241771/NA19913.final.cram

ftp://ftp.sra.ebi.ac.uk/vol1/run/ERR324/ERR3241773/NA20318.final.cram

ftp://ftp.sra.ebi.ac.uk/vol1/run/ERR324/ERR3241777/NA20362.final.cram

ftp://ftp.sra.ebi.ac.uk/vol1/run/ERR324/ERR3241775/NA20321.final.cram

ftp://ftp.sra.ebi.ac.uk/vol1/run/ERR324/ERR3241772/NA20274.final.cram

ftp://ftp.sra.ebi.ac.uk/vol1/run/ERR324/ERR3241774/NA20320.final.cram

ftp://ftp.sra.ebi.ac.uk/vol1/run/ERR398/ERR3989427/NA19702.final.cram

ftp://ftp.sra.ebi.ac.uk/vol1/run/ERR398/ERR3989449/NA19836.final.cram

ftp://ftp.sra.ebi.ac.uk/vol1/run/ERR398/ERR3989451/NA19918.final.cram

ftp://ftp.sra.ebi.ac.uk/vol1/run/ERR398/ERR3989456/NA20129.final.cram

ftp://ftp.sra.ebi.ac.uk/vol1/run/ERR398/ERR3989458/NA20358.final.cram

ftp://ftp.sra.ebi.ac.uk/vol1/run/ERR398/ERR3989448/NA19828.final.cram

ftp://ftp.sra.ebi.ac.uk/vol1/run/ERR398/ERR3989450/NA19902.final.cram

ftp://ftp.sra.ebi.ac.uk/vol1/run/ERR398/ERR3989455/NA20128.final.cram

ftp://ftp.sra.ebi.ac.uk/vol1/run/ERR398/ERR3989457/NA20279.final.cram

ftp://ftp.sra.ebi.ac.uk/vol1/run/ERR323/ERR3239915/NA19703.final.cram

ftp://ftp.sra.ebi.ac.uk/vol1/run/ERR323/ERR3239959/NA19818.final.cram

ftp://ftp.sra.ebi.ac.uk/vol1/run/ERR323/ERR3239961/NA19834.final.cram

ftp://ftp.sra.ebi.ac.uk/vol1/run/ERR323/ERR3239964/NA19901.final.cram

ftp://ftp.sra.ebi.ac.uk/vol1/run/ERR323/ERR3239969/NA19916.final.cram

ftp://ftp.sra.ebi.ac.uk/vol1/run/ERR323/ERR3239976/NA20276.final.cram

ftp://ftp.sra.ebi.ac.uk/vol1/run/ERR323/ERR3239981/NA20289.final.cram

ftp://ftp.sra.ebi.ac.uk/vol1/run/ERR323/ERR3239983/NA20294.final.cram

ftp://ftp.sra.ebi.ac.uk/vol1/run/ERR323/ERR3239988/NA20332.final.cram

ftp://ftp.sra.ebi.ac.uk/vol1/run/ERR323/ERR3239990/NA20340.final.cram

ftp://ftp.sra.ebi.ac.uk/vol1/run/ERR323/ERR3239995/NA20357.final.cram

ftp://ftp.sra.ebi.ac.uk/vol1/run/ERR323/ERR3239916/NA19704.final.cram

ftp://ftp.sra.ebi.ac.uk/vol1/run/ERR323/ERR3239960/NA19819.final.cram

ftp://ftp.sra.ebi.ac.uk/vol1/run/ERR323/ERR3239962/NA19835.final.cram

ftp://ftp.sra.ebi.ac.uk/vol1/run/ERR323/ERR3239965/NA19904.final.cram

ftp://ftp.sra.ebi.ac.uk/vol1/run/ERR323/ERR3239970/NA19917.final.cram

ftp://ftp.sra.ebi.ac.uk/vol1/run/ERR323/ERR3239977/NA20278.final.cram

ftp://ftp.sra.ebi.ac.uk/vol1/run/ERR323/ERR3239982/NA20291.final.cram

ftp://ftp.sra.ebi.ac.uk/vol1/run/ERR323/ERR3239984/NA20296.final.cram

ftp://ftp.sra.ebi.ac.uk/vol1/run/ERR324/ERR3240097/NA19984.final.cram

ftp://ftp.sra.ebi.ac.uk/vol1/run/ERR324/ERR3240099/NA20339.final.cram

ftp://ftp.sra.ebi.ac.uk/vol1/run/ERR323/ERR3239991/NA20342.final.cram

ftp://ftp.sra.ebi.ac.uk/vol1/run/ERR323/ERR3239996/NA20359.final.cram

ftp://ftp.sra.ebi.ac.uk/vol1/run/ERR324/ERR3240098/NA20298.final.cram

ftp://ftp.sra.ebi.ac.uk/vol1/run/ERR324/ERR3240100/NA20351.final.cram

ftp://ftp.sra.ebi.ac.uk/vol1/run/ERR324/ERR3241776/NA20355.final.cram

ftp://ftp.sra.ebi.ac.uk/vol1/run/ERR398/ERR3989428/NA19705.final.cram

ftp://ftp.sra.ebi.ac.uk/vol1/run/ERR398/ERR3989452/NA19919.final.cram

ftp://ftp.sra.ebi.ac.uk/vol1/run/ERR398/ERR3989453/NA19924.final.cram

ftp://ftp.sra.ebi.ac.uk/vol1/run/ERR398/ERR3989454/NA19983.final.cram

**Individuals with 30X WGS data in the 1000GenomeProject Utah residents (CEPH) with Northern and Western European ancestry group (CEU)**

ftp://ftp.sra.ebi.ac.uk/vol1/run/ERR323/ERR3239480/NA12718.final.cram

ftp://ftp.sra.ebi.ac.uk/vol1/run/ERR323/ERR3239482/NA12775.final.cram

ftp://ftp.sra.ebi.ac.uk/vol1/run/ERR323/ERR3239487/NA12842.final.cram

ftp://ftp.sra.ebi.ac.uk/vol1/run/ERR323/ERR3239280/NA07037.final.cram

ftp://ftp.sra.ebi.ac.uk/vol1/run/ERR323/ERR3239286/NA11829.final.cram

ftp://ftp.sra.ebi.ac.uk/vol1/run/ERR323/ERR3239293/NA11918.final.cram

ftp://ftp.sra.ebi.ac.uk/vol1/run/ERR323/ERR3239298/NA11994.final.cram

ftp://ftp.sra.ebi.ac.uk/vol1/run/ERR323/ERR3239301/NA12004.final.cram

ftp://ftp.sra.ebi.ac.uk/vol1/run/ERR323/ERR3239643/NA11932.final.cram

ftp://ftp.sra.ebi.ac.uk/vol1/run/ERR323/ERR3239307/NA12144.final.cram

ftp://ftp.sra.ebi.ac.uk/vol1/run/ERR323/ERR3239312/NA12249.final.cram

ftp://ftp.sra.ebi.ac.uk/vol1/run/ERR323/ERR3239319/NA12750.final.cram

ftp://ftp.sra.ebi.ac.uk/vol1/run/ERR323/ERR3239324/NA12763.final.cram

ftp://ftp.sra.ebi.ac.uk/vol1/run/ERR323/ERR3239326/NA12812.final.cram

ftp://ftp.sra.ebi.ac.uk/vol1/run/ERR323/ERR3239333/NA12874.final.cram

ftp://ftp.sra.ebi.ac.uk/vol1/run/ERR323/ERR3239461/NA11892.final.cram

ftp://ftp.sra.ebi.ac.uk/vol1/run/ERR323/ERR3239466/NA12273.final.cram

ftp://ftp.sra.ebi.ac.uk/vol1/run/ERR323/ERR3239468/NA12282.final.cram

ftp://ftp.sra.ebi.ac.uk/vol1/run/ERR323/ERR3239473/NA12342.final.cram

ftp://ftp.sra.ebi.ac.uk/vol1/run/ERR323/ERR3239484/NA12778.final.cram

ftp://ftp.sra.ebi.ac.uk/vol1/run/ERR323/ERR3239489/NA12889.final.cram

ftp://ftp.sra.ebi.ac.uk/vol1/run/ERR323/ERR3239277/NA06986.final.cram

ftp://ftp.sra.ebi.ac.uk/vol1/run/ERR323/ERR3239481/NA12748.final.cram

ftp://ftp.sra.ebi.ac.uk/vol1/run/ERR323/ERR3239483/NA12777.final.cram

ftp://ftp.sra.ebi.ac.uk/vol1/run/ERR323/ERR3239488/NA12843.final.cram

ftp://ftp.sra.ebi.ac.uk/vol1/run/ERR323/ERR3239283/NA07357.final.cram

ftp://ftp.sra.ebi.ac.uk/vol1/run/ERR323/ERR3239288/NA11831.final.cram

ftp://ftp.sra.ebi.ac.uk/vol1/run/ERR323/ERR3239290/NA11840.final.cram

ftp://ftp.sra.ebi.ac.uk/vol1/run/ERR323/ERR3239295/NA11920.final.cram

ftp://ftp.sra.ebi.ac.uk/vol1/run/ERR323/ERR3239303/NA12006.final.cram

ftp://ftp.sra.ebi.ac.uk/vol1/run/ERR323/ERR3239645/NA12046.final.cram

ftp://ftp.sra.ebi.ac.uk/vol1/run/ERR323/ERR3239309/NA12155.final.cram

ftp://ftp.sra.ebi.ac.uk/vol1/run/ERR323/ERR3239314/NA12414.final.cram

ftp://ftp.sra.ebi.ac.uk/vol1/run/ERR323/ERR3239316/NA12716.final.cram

ftp://ftp.sra.ebi.ac.uk/vol1/run/ERR323/ERR3239321/NA12760.final.cram

ftp://ftp.sra.ebi.ac.uk/vol1/run/ERR323/ERR3239328/NA12814.final.cram

ftp://ftp.sra.ebi.ac.uk/vol1/run/ERR323/ERR3239276/NA06985.final.cram

ftp://ftp.sra.ebi.ac.uk/vol1/run/ERR323/ERR3239281/NA07051.final.cram

ftp://ftp.sra.ebi.ac.uk/vol1/run/ERR323/ERR3239282/NA07347.final.cram

ftp://ftp.sra.ebi.ac.uk/vol1/run/ERR323/ERR3239287/NA11830.final.cram

ftp://ftp.sra.ebi.ac.uk/vol1/run/ERR323/ERR3239289/NA11832.final.cram

ftp://ftp.sra.ebi.ac.uk/vol1/run/ERR323/ERR3239294/NA11919.final.cram

ftp://ftp.sra.ebi.ac.uk/vol1/run/ERR323/ERR3239302/NA12005.final.cram

ftp://ftp.sra.ebi.ac.uk/vol1/run/ERR323/ERR3239644/NA11933.final.cram

ftp://ftp.sra.ebi.ac.uk/vol1/run/ERR323/ERR3239308/NA12154.final.cram

ftp://ftp.sra.ebi.ac.uk/vol1/run/ERR323/ERR3239313/NA12287.final.cram

ftp://ftp.sra.ebi.ac.uk/vol1/run/ERR323/ERR3239315/NA12489.final.cram

ftp://ftp.sra.ebi.ac.uk/vol1/run/ERR323/ERR3239320/NA12751.final.cram

ftp://ftp.sra.ebi.ac.uk/vol1/run/ERR323/ERR3239456/NA07056.final.cram

ftp://ftp.sra.ebi.ac.uk/vol1/run/ERR323/ERR3239327/NA12813.final.cram

ftp://ftp.sra.ebi.ac.uk/vol1/run/ERR323/ERR3239334/NA12878.final.cram

ftp://ftp.sra.ebi.ac.uk/vol1/run/ERR323/ERR3239458/NA06984.final.cram

ftp://ftp.sra.ebi.ac.uk/vol1/run/ERR323/ERR3239463/NA11930.final.cram

ftp://ftp.sra.ebi.ac.uk/vol1/run/ERR323/ERR3239470/NA12286.final.cram

ftp://ftp.sra.ebi.ac.uk/vol1/run/ERR323/ERR3239475/NA12348.final.cram

ftp://ftp.sra.ebi.ac.uk/vol1/run/ERR323/ERR3239477/NA12399.final.cram

ftp://ftp.sra.ebi.ac.uk/vol1/run/ERR323/ERR3239462/NA11893.final.cram

ftp://ftp.sra.ebi.ac.uk/vol1/run/ERR323/ERR3239469/NA12283.final.cram

ftp://ftp.sra.ebi.ac.uk/vol1/run/ERR323/ERR3239474/NA12347.final.cram

ftp://ftp.sra.ebi.ac.uk/vol1/run/ERR398/ERR3989266/NA06997.final.cram

ftp://ftp.sra.ebi.ac.uk/vol1/run/ERR398/ERR3989271/NA07031.final.cram

ftp://ftp.sra.ebi.ac.uk/vol1/run/ERR398/ERR3989273/NA07045.final.cram

ftp://ftp.sra.ebi.ac.uk/vol1/run/ERR398/ERR3989278/NA07349.final.cram

ftp://ftp.sra.ebi.ac.uk/vol1/run/ERR398/ERR3989282/NA10835.final.cram

ftp://ftp.sra.ebi.ac.uk/vol1/run/ERR398/ERR3989287/NA10840.final.cram

ftp://ftp.sra.ebi.ac.uk/vol1/run/ERR398/ERR3989294/NA10855.final.cram

ftp://ftp.sra.ebi.ac.uk/vol1/run/ERR398/ERR3989299/NA10861.final.cram

ftp://ftp.sra.ebi.ac.uk/vol1/run/ERR398/ERR3989304/NA11882.final.cram

ftp://ftp.sra.ebi.ac.uk/vol1/run/ERR398/ERR3989309/NA12057.final.cram

ftp://ftp.sra.ebi.ac.uk/vol1/run/ERR398/ERR3989311/NA12146.final.cram

ftp://ftp.sra.ebi.ac.uk/vol1/run/ERR398/ERR3989316/NA12274.final.cram

ftp://ftp.sra.ebi.ac.uk/vol1/run/ERR398/ERR3989323/NA12386.final.cram

ftp://ftp.sra.ebi.ac.uk/vol1/run/ERR398/ERR3989326/NA12739.final.cram

ftp://ftp.sra.ebi.ac.uk/vol1/run/ERR398/ERR3989334/NA12817.final.cram

ftp://ftp.sra.ebi.ac.uk/vol1/run/ERR398/ERR3989339/NA12875.final.cram

ftp://ftp.sra.ebi.ac.uk/vol1/run/ERR398/ERR3989341/NA12891.final.cram

ftp://ftp.sra.ebi.ac.uk/vol1/run/ERR398/ERR3989263/NA06991.final.cram

ftp://ftp.sra.ebi.ac.uk/vol1/run/ERR398/ERR3989268/NA07019.final.cram

ftp://ftp.sra.ebi.ac.uk/vol1/run/ERR398/ERR3989275/NA07345.final.cram

ftp://ftp.sra.ebi.ac.uk/vol1/run/ERR398/ERR3989280/NA10830.final.cram

ftp://ftp.sra.ebi.ac.uk/vol1/run/ERR398/ERR3989284/NA10837.final.cram

ftp://ftp.sra.ebi.ac.uk/vol1/run/ERR398/ERR3989289/NA10843.final.cram

ftp://ftp.sra.ebi.ac.uk/vol1/run/ERR398/ERR3989296/NA10857.final.cram

ftp://ftp.sra.ebi.ac.uk/vol1/run/ERR398/ERR3989301/NA10864.final.cram

ftp://ftp.sra.ebi.ac.uk/vol1/run/ERR398/ERR3989306/NA11917.final.cram

ftp://ftp.sra.ebi.ac.uk/vol1/run/ERR398/ERR3989313/NA12239.final.cram

ftp://ftp.sra.ebi.ac.uk/vol1/run/ERR398/ERR3989318/NA12335.final.cram

ftp://ftp.sra.ebi.ac.uk/vol1/run/ERR398/ERR3989328/NA12752.final.cram

ftp://ftp.sra.ebi.ac.uk/vol1/run/ERR398/ERR3989331/NA12767.final.cram

ftp://ftp.sra.ebi.ac.uk/vol1/run/ERR398/ERR3989336/NA12832.final.cram

ftp://ftp.sra.ebi.ac.uk/vol1/run/ERR398/ERR3989267/NA07014.final.cram

ftp://ftp.sra.ebi.ac.uk/vol1/run/ERR398/ERR3989272/NA07034.final.cram

ftp://ftp.sra.ebi.ac.uk/vol1/run/ERR398/ERR3989274/NA07055.final.cram

ftp://ftp.sra.ebi.ac.uk/vol1/run/ERR398/ERR3989279/NA07435.final.cram

ftp://ftp.sra.ebi.ac.uk/vol1/run/ERR398/ERR3989283/NA10836.final.cram

ftp://ftp.sra.ebi.ac.uk/vol1/run/ERR398/ERR3989288/NA10842.final.cram

ftp://ftp.sra.ebi.ac.uk/vol1/run/ERR398/ERR3989295/NA10856.final.cram

ftp://ftp.sra.ebi.ac.uk/vol1/run/ERR398/ERR3989300/NA10863.final.cram

ftp://ftp.sra.ebi.ac.uk/vol1/run/ERR398/ERR3989302/NA10865.final.cram

ftp://ftp.sra.ebi.ac.uk/vol1/run/ERR398/ERR3989305/NA11891.final.cram

ftp://ftp.sra.ebi.ac.uk/vol1/run/ERR398/ERR3989310/NA12145.final.cram

ftp://ftp.sra.ebi.ac.uk/vol1/run/ERR398/ERR3989312/NA12236.final.cram

ftp://ftp.sra.ebi.ac.uk/vol1/run/ERR398/ERR3989327/NA12740.final.cram

ftp://ftp.sra.ebi.ac.uk/vol1/run/ERR398/ERR3989330/NA12766.final.cram

ftp://ftp.sra.ebi.ac.uk/vol1/run/ERR398/ERR3989335/NA12818.final.cram

ftp://ftp.sra.ebi.ac.uk/vol1/run/ERR398/ERR3989342/NA12892.final.cram

ftp://ftp.sra.ebi.ac.uk/vol1/run/ERR323/ERR3239485/NA12827.final.cram

ftp://ftp.sra.ebi.ac.uk/vol1/run/ERR323/ERR3239490/NA12890.final.cram

ftp://ftp.sra.ebi.ac.uk/vol1/run/ERR323/ERR3239278/NA06994.final.cram

ftp://ftp.sra.ebi.ac.uk/vol1/run/ERR323/ERR3239486/NA12829.final.cram

ftp://ftp.sra.ebi.ac.uk/vol1/run/ERR323/ERR3239284/NA10847.final.cram

ftp://ftp.sra.ebi.ac.uk/vol1/run/ERR323/ERR3239291/NA11881.final.cram

ftp://ftp.sra.ebi.ac.uk/vol1/run/ERR323/ERR3239296/NA11931.final.cram

ftp://ftp.sra.ebi.ac.uk/vol1/run/ERR323/ERR3239299/NA11995.final.cram

ftp://ftp.sra.ebi.ac.uk/vol1/run/ERR323/ERR3239305/NA12044.final.cram

ftp://ftp.sra.ebi.ac.uk/vol1/run/ERR323/ERR3239310/NA12156.final.cram

ftp://ftp.sra.ebi.ac.uk/vol1/run/ERR323/ERR3239317/NA12717.final.cram

ftp://ftp.sra.ebi.ac.uk/vol1/run/ERR323/ERR3239322/NA12761.final.cram

ftp://ftp.sra.ebi.ac.uk/vol1/run/ERR323/ERR3239329/NA12815.final.cram

ftp://ftp.sra.ebi.ac.uk/vol1/run/ERR323/ERR3239331/NA12872.final.cram

ftp://ftp.sra.ebi.ac.uk/vol1/run/ERR323/ERR3239279/NA07000.final.cram

ftp://ftp.sra.ebi.ac.uk/vol1/run/ERR323/ERR3239285/NA10851.final.cram

ftp://ftp.sra.ebi.ac.uk/vol1/run/ERR323/ERR3239292/NA11894.final.cram

ftp://ftp.sra.ebi.ac.uk/vol1/run/ERR323/ERR3239297/NA11992.final.cram

ftp://ftp.sra.ebi.ac.uk/vol1/run/ERR323/ERR3239300/NA12003.final.cram

ftp://ftp.sra.ebi.ac.uk/vol1/run/ERR323/ERR3239642/NA07048.final.cram

ftp://ftp.sra.ebi.ac.uk/vol1/run/ERR323/ERR3239304/NA12043.final.cram

ftp://ftp.sra.ebi.ac.uk/vol1/run/ERR323/ERR3239306/NA12045.final.cram

ftp://ftp.sra.ebi.ac.uk/vol1/run/ERR323/ERR3239311/NA12234.final.cram

ftp://ftp.sra.ebi.ac.uk/vol1/run/ERR323/ERR3239318/NA12749.final.cram

ftp://ftp.sra.ebi.ac.uk/vol1/run/ERR323/ERR3239323/NA12762.final.cram

ftp://ftp.sra.ebi.ac.uk/vol1/run/ERR323/ERR3239325/NA12776.final.cram

ftp://ftp.sra.ebi.ac.uk/vol1/run/ERR323/ERR3239330/NA12828.final.cram

ftp://ftp.sra.ebi.ac.uk/vol1/run/ERR323/ERR3239332/NA12873.final.cram

ftp://ftp.sra.ebi.ac.uk/vol1/run/ERR323/ERR3239459/NA06989.final.cram

ftp://ftp.sra.ebi.ac.uk/vol1/run/ERR323/ERR3239464/NA12058.final.cram

ftp://ftp.sra.ebi.ac.uk/vol1/run/ERR323/ERR3239471/NA12340.final.cram

ftp://ftp.sra.ebi.ac.uk/vol1/run/ERR323/ERR3239476/NA12383.final.cram

ftp://ftp.sra.ebi.ac.uk/vol1/run/ERR323/ERR3239478/NA12400.final.cram

ftp://ftp.sra.ebi.ac.uk/vol1/run/ERR323/ERR3239460/NA11843.final.cram

ftp://ftp.sra.ebi.ac.uk/vol1/run/ERR323/ERR3239465/NA12272.final.cram

ftp://ftp.sra.ebi.ac.uk/vol1/run/ERR323/ERR3239467/NA12275.final.cram

ftp://ftp.sra.ebi.ac.uk/vol1/run/ERR323/ERR3239472/NA12341.final.cram

ftp://ftp.sra.ebi.ac.uk/vol1/run/ERR323/ERR3239479/NA12413.final.cram

ftp://ftp.sra.ebi.ac.uk/vol1/run/ERR324/ERR3243162/NA12830.final.cram

ftp://ftp.sra.ebi.ac.uk/vol1/run/ERR324/ERR3243163/NA12546.final.cram

ftp://ftp.sra.ebi.ac.uk/vol1/run/ERR398/ERR3989264/NA06993.final.cram

ftp://ftp.sra.ebi.ac.uk/vol1/run/ERR398/ERR3989269/NA07022.final.cram

ftp://ftp.sra.ebi.ac.uk/vol1/run/ERR398/ERR3989276/NA07346.final.cram

ftp://ftp.sra.ebi.ac.uk/vol1/run/ERR398/ERR3989281/NA10831.final.cram

ftp://ftp.sra.ebi.ac.uk/vol1/run/ERR398/ERR3989285/NA10838.final.cram

ftp://ftp.sra.ebi.ac.uk/vol1/run/ERR398/ERR3989290/NA10845.final.cram

ftp://ftp.sra.ebi.ac.uk/vol1/run/ERR398/ERR3989292/NA10852.final.cram

ftp://ftp.sra.ebi.ac.uk/vol1/run/ERR398/ERR3989297/NA10859.final.cram

ftp://ftp.sra.ebi.ac.uk/vol1/run/ERR398/ERR3989307/NA11993.final.cram

ftp://ftp.sra.ebi.ac.uk/vol1/run/ERR398/ERR3989314/NA12248.final.cram

ftp://ftp.sra.ebi.ac.uk/vol1/run/ERR398/ERR3989319/NA12336.final.cram

ftp://ftp.sra.ebi.ac.uk/vol1/run/ERR398/ERR3989321/NA12344.final.cram

ftp://ftp.sra.ebi.ac.uk/vol1/run/ERR398/ERR3989324/NA12485.final.cram

ftp://ftp.sra.ebi.ac.uk/vol1/run/ERR398/ERR3989329/NA12753.final.cram

ftp://ftp.sra.ebi.ac.uk/vol1/run/ERR398/ERR3989332/NA12801.final.cram

ftp://ftp.sra.ebi.ac.uk/vol1/run/ERR398/ERR3989337/NA12864.final.cram

ftp://ftp.sra.ebi.ac.uk/vol1/run/ERR324/ERR3243162/NA12830.final.cram

ftp://ftp.sra.ebi.ac.uk/vol1/run/ERR324/ERR3243163/NA12546.final.cram

ftp://ftp.sra.ebi.ac.uk/vol1/run/ERR398/ERR3989264/NA06993.final.cram

ftp://ftp.sra.ebi.ac.uk/vol1/run/ERR398/ERR3989269/NA07022.final.cram

ftp://ftp.sra.ebi.ac.uk/vol1/run/ERR398/ERR3989276/NA07346.final.cram

ftp://ftp.sra.ebi.ac.uk/vol1/run/ERR398/ERR3989281/NA10831.final.cram

ftp://ftp.sra.ebi.ac.uk/vol1/run/ERR398/ERR3989285/NA10838.final.cram

ftp://ftp.sra.ebi.ac.uk/vol1/run/ERR398/ERR3989290/NA10845.final.cram

ftp://ftp.sra.ebi.ac.uk/vol1/run/ERR398/ERR3989292/NA10852.final.cram

ftp://ftp.sra.ebi.ac.uk/vol1/run/ERR398/ERR3989297/NA10859.final.cram

ftp://ftp.sra.ebi.ac.uk/vol1/run/ERR398/ERR3989307/NA11993.final.cram

ftp://ftp.sra.ebi.ac.uk/vol1/run/ERR398/ERR3989314/NA12248.final.cram

ftp://ftp.sra.ebi.ac.uk/vol1/run/ERR398/ERR3989319/NA12336.final.cram

ftp://ftp.sra.ebi.ac.uk/vol1/run/ERR398/ERR3989321/NA12344.final.cram

ftp://ftp.sra.ebi.ac.uk/vol1/run/ERR398/ERR3989324/NA12485.final.cram

ftp://ftp.sra.ebi.ac.uk/vol1/run/ERR398/ERR3989329/NA12753.final.cram
